# Developmental Dysfunction of VIP Interneurons Impairs Cortical Circuits

**DOI:** 10.1101/077891

**Authors:** Renata Batista-Brito, Martin Vinck, Katie A. Ferguson, David Laubender, Gyorgy Lur, Michael J. Higley, Jessica A. Cardin

## Abstract

Current evidence suggests that dysregulation of GABAergic interneurons contributes to neural and behavioral deficits in several neurodevelopmental disorders, including schizophrenia. However, there are multiple populations of interneurons and their respective roles in psychiatric disease remain poorly explored. Neuregulin 1 (*NRG1*) and its interneuron-specific tyrosine kinase receptor *ERBB4* are risk genes for schizophrenia, and the Nrg1/ErbB4 pathway is important for normal cortical development. Using a conditional ErbB4 deletion model, we directly tested the role of vasoactive intestinal peptide (VIP)-expressing interneurons in schizophrenia-related deficits *in vivo*. ErbB4 removal from VIP interneurons during development leads to changes in their activity, along with severe dysregulation of the temporal organization and state-dependence of cortical activity. As a result of these neural circuit alterations, animals in which VIP interneurons lack ErbB4 exhibit behavioral abnormalities, reduced cortical responses to sensory stimuli, and impaired sensory learning. Our data support a key role for VIP interneurons in normal cortical circuit development and suggest that their disruption contributes to pathophysiology in the ErbB4 model of schizophrenia. These findings provide a new perspective on the role of GABAergic interneuron diversity in the disruption of cortical function in complex psychiatric diseases.

## INTRODUCTION

Dysregulation of GABAergic inhibition in the brain is a candidate mechanism underlying schizophrenia and other neurodevelopmental disorders (Uhlhaas and Singer, 2010). Schizophrenic patients exhibit altered gamma oscillations and high co-morbidity for epilepsy, two forms of patterned brain activity regulated by GABAergic signaling (Karouni et al., 2010; Lee et al., 2010b; Uhlhaas and Singer, 2010). Initial symptoms are exhibited during adolescence and young adulthood, the time when GABAergic circuits are maturing (Fishell and Rudy, 2010). However, relatively little is known about the precise contribution of developmental GABAergic cellular dysfunction to subsequent altered neural network activity and cognitive and perceptual impairments in schizophrenia or other complex psychiatric disorders.

Cortical tissue from schizophrenic patients shows decreased expression of markers for multiple distinct populations of interneurons, including those that co-express parvalbumin (PV), somatostatin (SST), and vasoactive intestinal peptide (VIP) (Hashimoto et al., 2008; Fung et al., 2010; Volk et al., 2012; Fung et al., 2014; Joshi et al., 2014). The respective contributions of these diverse interneuron populations to disease-related deficits are unknown. Several studies have focused on a role for PV interneurons, whose function may contribute to disease-related alterations in neural activity patterns and behavior (Hikida et al., 2007; Fazzari et al., 2010; Wen et al., 2010). However, VIP-expressing interneurons (VIP-INs) have recently gained attention as important regulators of cortical function (Lee et al., 2013; Pi et al., 2013; Fu et al., 2014; Karnani et al., 2016). VIP-INs are regular-spiking, dendrite-targeting cells preferentially found in superficial cortical layers. They receive local and long-range excitatory inputs as well as serotonergic and cholinergic afferents, two neuromodulatory systems affected in schizophrenia and other psychiatric diseases (Gray and Roth, 2007; Lee et al., 2010a; Higley and Picciotto, 2014; Pronneke et al., 2015; Wall et al., 2016). VIP-INs are strongly recruited by negative or noxious stimuli and arousing events such as the onset of motor activity (Lee et al., 2013; Pi et al., 2013; Fu et al., 2014). In turn, they regulate cortical excitatory activity and sensory response gain through inhibition of pyramidal neurons and other interneurons (Lee et al., 2013; Pfeffer et al., 2013; Pi et al., 2013; Fu et al., 2014). VIP-INs integrate into cortical circuits early in postnatal life (Miyoshi et al., 2015) However, despite their powerful influence on cortical activity, nothing is known about how they shape normal development of the cerebral cortex or whether they contribute to neural and cognitive deficits in neurodevelopmental disorders such as schizophrenia.

The signaling factor Neuregulin-1 (Nrg-1) and its membrane-bound tyrosine kinase receptor ERBB4 (ErbB4) are elements of a signaling pathway critical for the proper development of cortical and hippocampal circuits. ErbB4 expression is restricted to GABAergic neurons in the cortex (Yau et al., 2003; Flames et al., 2004; Neddens et al., 2011), and this pathway plays a role in regulating GABAergic synaptic development (Fazzari et al., 2010; Del Pino et al., 2013). Nrg-1 and ErbB4 are strong candidate genes for schizophrenia (Rico and Marin, 2011; Mei and Nave, 2014). Mice globally lacking Nrg-1 or ErbB4 exhibit key behavioral deficits associated with this neurodevelopmental disease, including impaired working and spatial memory, hyperactivity, decreased prepulse inhibition, and disrupted social and emotional behaviors (Karl et al., 2007; Rico and Marin, 2011; Shamir et al., 2012). ErbB4 signaling is necessary for normal interneuron migration and synapse formation during development (Fazzari et al., 2010; Shamir et al., 2012), and global disruptions of the Nrg-1/ErbB4 pathway lead to decreased GABA release in the mature cerebral cortex (Woo et al., 2007; Mei and Xiong, 2008; Neddens and Buonanno, 2010; Ting et al., 2011) and impaired circuit function in mature animals (Barz et al., 2015, 2016). Disruption of the Nrg1-ErbB4 pathway specifically in PV interneurons reduces synaptic input to these cells and appears to contribute to the behavioral and neural phenotypes observed in the total ErbB4 deletion model (Chen et al., 2010; Wen et al., 2010; Shamir et al., 2012; Del Pino et al., 2013). However, the impact of Nrg-1/ErbB4 signaling in other cortical interneuron populations, such as VIP cells, is unknown.

Here we tested the influence of ErbB4 signaling in VIP-INs on the development and function of cortical circuits. We find that deletion of ErbB4 from VIP-INs causes a profound dysregulation of cortical activity that appears restricted to excitatory neurons. VIP-IN-specific loss of ErbB4 also perturbs behavioral state-dependent regulation of cortical circuits and impairs sensory processing. Finally, animals lacking ErbB4 in VIP-INs exhibit impairments in sensory learning and other behaviors, reminiscent of dysfunction seen in both global ErbB4 deletion and schizophrenic patients, suggesting a potential causal role for these interneurons in disease pathophysiology.

## RESULTS

### Developmental deletion of ErbB4 from VIP interneurons

We directly tested whether developmental dysfunction of VIP interneurons contributes to disease-related cortical deficits following disruption of the ErbB4-Nrg1 signaling pathway, using mouse primary visual cortex (V1) as a model for local circuit function. We found that ErbB4 expression in V1 was restricted to GABAergic interneurons and was present in most VIP- and parvalbumin- (PV) and some somatostatin-expressing (SST) interneurons (Figures 1A-B, S1). We developmentally ablated Nrg1-ErbB4 signaling specifically in VIP-INs early in postnatal life (Miyoshi et al., 2015) by generating ErbB4^F/F^,VIP^Cre^ mice (Fig. 1A-B). We did not observe a decrease in the overall density of VIP-INs in the mutants (Figure 1C), suggesting that ErbB4 deletion did not lead to death or altered migration of these cells. To examine the broad functional consequences of ErbB4 deletion from VIP-INs, we performed a series of behavioral assays disrupted in models of schizophrenia. Similarly to ErbB4 null mice (Shamir et al., 2012), ErbB4^F/F^,VIP^Cre^ mutants exhibited hyperactivity and reduced levels of anxiety as shown by open field and marble burying behavior assays (Figure S2), suggesting that VIP interneuron dysregulation contributes to disease-related phenotypes.

**Figure 1.**
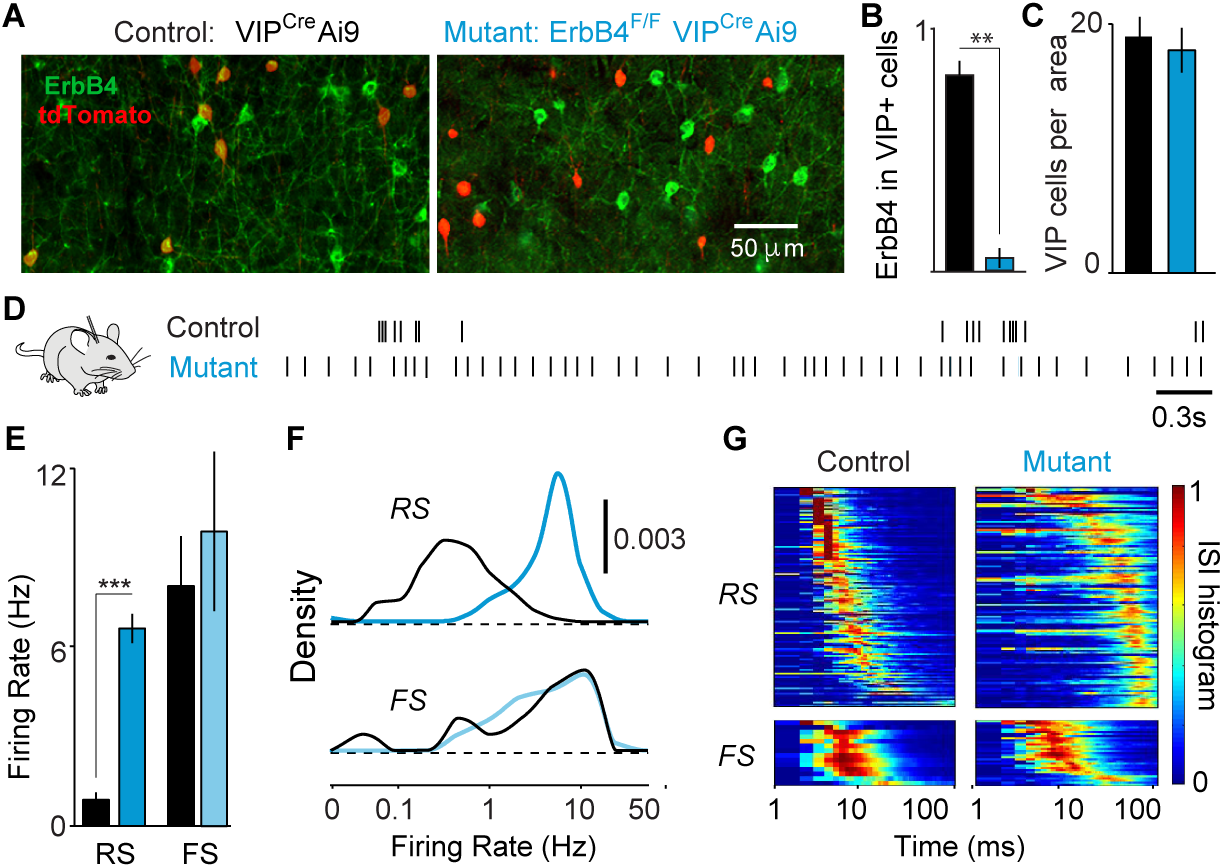
ErbB4 deletion from VIP interneurons alters cortical spiking. (A) ErbB4 and tdTomato immunohistochemistry in V1 cortex of control and mutant mice. (B) ErbB4 expression in VIP interneurons in controls (black) (n=6 mice) and mutants (cyan; n=6). (C) Number of VIP fate-mapped cells per optical area in control (black) and mutant (cyan) mice. (D) Spike trains of example RS cells. (E) Average firing rate during quiescence. Controls: 153 RS, 15 FS cells, 8 mice. Mutants: 134 RS, 32 FS cells, 8 mice. (F) Distribution of firing rates across population. (G) Inter-spike interval histograms (normalized to max) during quiescence for all cells. Error bars show s.e.m., p**<0.01, p***<0.001.

### Loss of key forms of temporally patterned cortical activity in ErbB4^F/F^,VIP^Cre^ mutants

Schizophrenia is associated with alterations in the balance of excitation and inhibition in cortical circuits (Uhlhaas and Singer, 2010). To assay the effects of ErbB4 deletion on neural activity, we performed extracellular recordings of regular spiking (RS; putative excitatory) neurons and fast spiking (FS; putative PV inhibitory) interneurons throughout cortical layers 2-6 in awake, head-fixed adult mutant and control mice (Figures 1, S3, S4, Experimental Procedures). VIP-INs are thought to regulate cortical circuits primarily by inhibiting other interneurons (Pfeffer et al., 2013; Pi et al., 2013; Fu et al., 2014), and we therefore predicted that mutants would exhibit both suppression of RS cells and changes in FS cell activity due to increased inhibition. Surprisingly, deletion of ErbB4 from VIP-INs led to marked increases in the spontaneous cortical activity patterns of RS, while FS cells were unchanged. RS cells in mutants exhibited ~4-fold higher spontaneous firing rates than those in controls (Figure 1D-F). In addition, RS cells in mutants displayed abnormal temporal spiking properties, including reduced bursting (Figure 1G, S4C-D) and decreased firing rate variability (Figure S4E). ErbB4 deletion from VIP interneurons was also associated with a broadening of RS action potentials (Figure S3B,D). These data suggest that early deletion of ErbB4 specifically from VIP-INs leads to hyperexcitability of excitatory neurons in the cortex.

Rhythmic neuronal synchronization is a core feature of healthy cortical activity, and reduced neural synchrony is a hallmark of schizophrenia (Uhlhaas and Singer, 2010; Gandal et al., 2012). To examine the impact of ErbB4 deletion in VIP interneurons on the temporal patterning of cortical activity, we simultaneously recorded spiking activity and local field potentials (LFPs) in V1 (Figure 2A). Despite showing only modest changes in LFP power (Figures 2A, S5A), mutants showed a near-complete reduction in the phase-locking of individual RS cells to low-frequency and gamma LFP oscillations regardless of behavioral state (Figure 2A-D, S5B). Correlations between the spiking of simultaneously recorded RS-RS and RS-FS pairs were abolished in mutants (Figures 2E, S5C), further indicating a loss of synchronous firing. FS cells typically exhibit a high level of synchrony with other FS cells, largely due to dense chemical and electrical synaptic connectivity (Gibson et al., 1999). Surprisingly, FS-FS synchrony was unaffected in mutants (Figure 2E). Together with the unchanged firing rates of FS cells in mutant animals (Figure 1), these findings suggest that the coordinated activity of RS and FS cell populations, a prominent element of cortical circuit function, is decoupled as a result of early VIP-IN dysfunction. Our data further indicate that early ErbB4 deletion from VIP-INs eliminates excitatory synchrony in cortical networks, disrupting several key forms of temporally organized activity that are important for information processing in cortical circuits (Fries, 2009).

**Figure 2.**
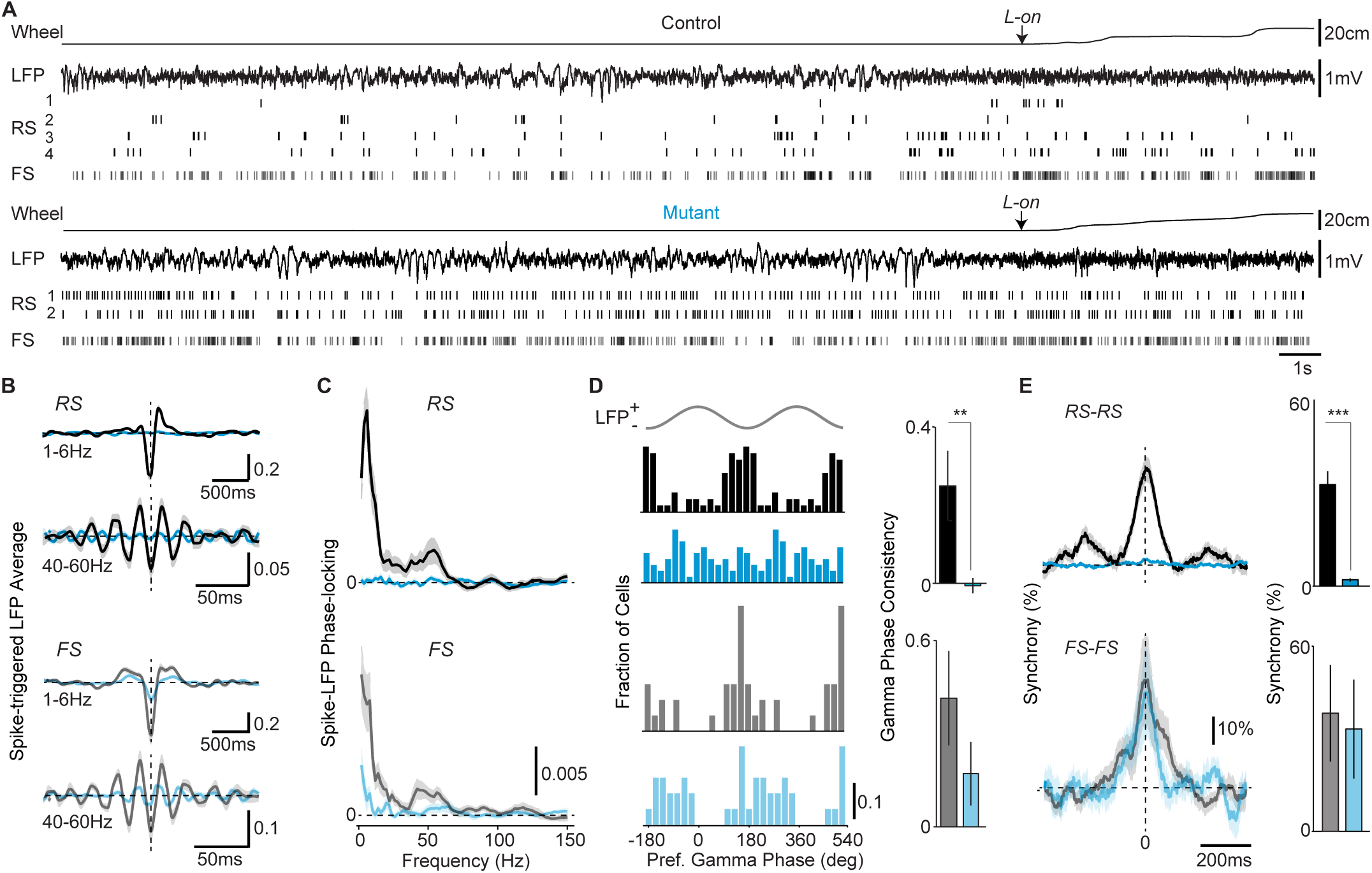
Loss of VIP ErbB4 disrupts the temporal organization of cortical activity. (A) Example wheel and LFP traces, with single-unit activity, around locomotion-onset (L-on). (B) Average spike-triggered LFP average in 40-60Hz and 1-6Hz bands during locomotion. Controls: 55 RS, 23 FS cells, 7 mice. Mutants: 61 RS, 23 FS, 8 mice. (C) Average spike-LFP phase-locking during locomotion. Control vs mutant significant in 1-6Hz (RS: p<0.05) and 40-60Hz (RS: p<0.05; FS: p<0.05). (D) Left: Preferred LFP gamma-phase of firing during locomotion. Right: Consistency of preferred LFP gamma-phases. (E) Left: Average normalized cross-correlograms during quiescence. Right: Percent-wise increase in zero-lag coincidences. RS-RS: Controls 192 pairs, 14 mice; Mutants 227 pairs, 5 mice. FS-FS: Controls n=15 pairs, 4 mice; Mutants 22 pairs, 3 mice. Error bars and shadings show s.e.m., p**<0.01, p***<0.001.

### ErbB4 deletion from VIP interneurons abolishes cortical state transitions

Because VIP-INs are thought to contribute to arousal-mediated changes in cortical activity during locomotion (Fu et al., 2014), we tested the impact of ErbB4 deletion from these cells on the ability of cortical circuits to follow transitions in behavioral state. We recorded extracellular signals in V1 cortex of mice transitioning between quiescent and active periods (Vinck et al., 2015). In contrast to robust increases in spiking in control animals, RS cells in mutants showed no significant change in firing at locomotion onset (Figure 3A-C). Likewise, RS cells showed no significant change at locomotion offset, a separate period of high global arousal that is independent of motor activity and normally associated with decreased firing rates (Figure 3B,D) (Vinck et al., 2015). These data indicate an extensive loss of the cortical response to behavioral arousal and suggest that ErbB4 deletion from VIP-INs prevents locomotion- and arousal-related signals from reaching cortical excitatory neurons.

**Figure 3.**
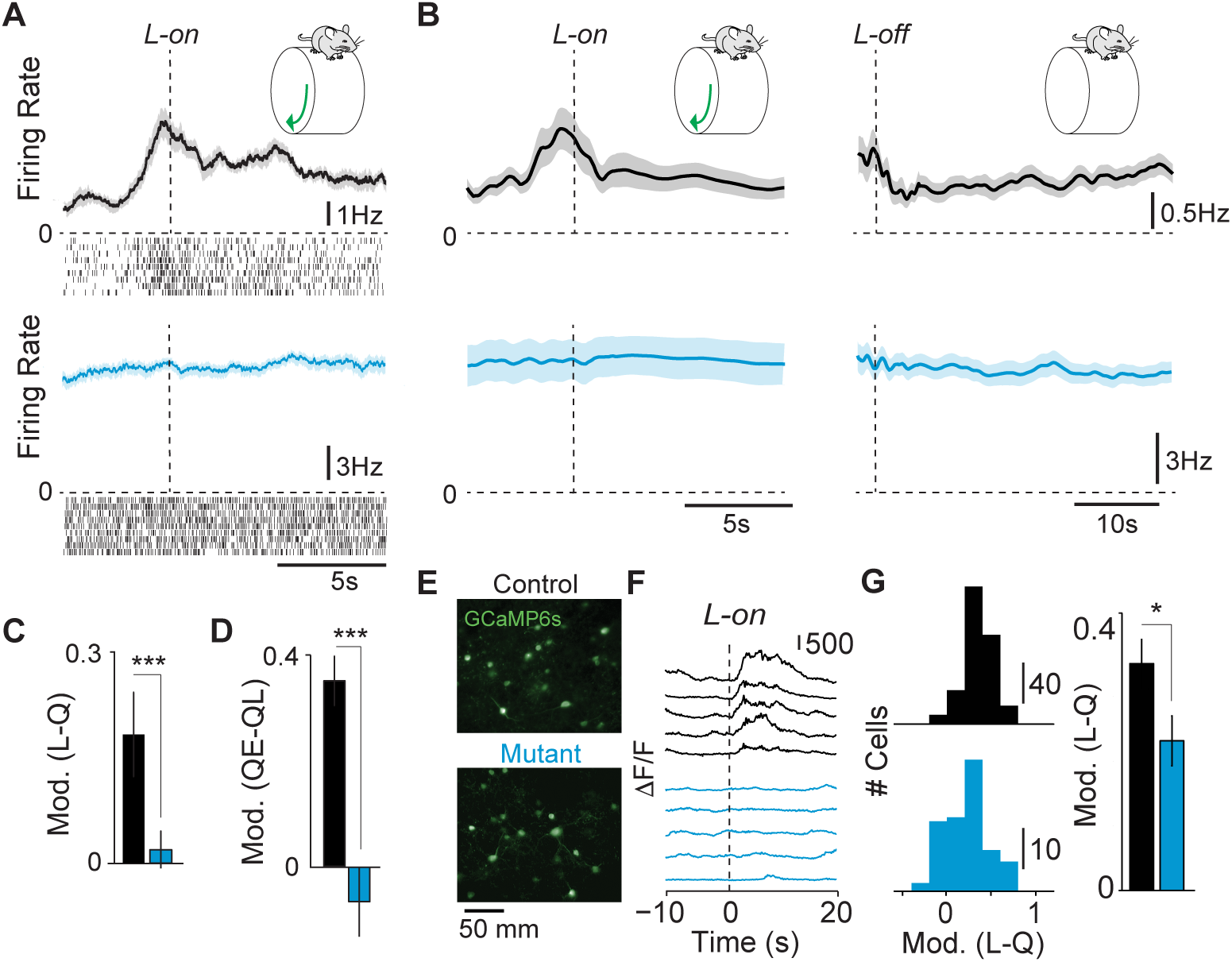
VIP ErbB4 deletion abolishes cortical state transitions. (A) Average firing rate and raster plots for example RS cells around locomotion onset (L-on) in control and mutant. (B) Population average change in RS firing rate around locomotion on- (left panel) and offset (right panel). Controls: 85 cells, 5 mice. Mutants: 72 cells, 8 mice. (C) Firing rate modulation in early locomotion period (L; -0.5 to 0.5s around L-on) as compared to quiescence (Q). (D) Firing rate modulation in late (QL) vs. early (QE) quiescence. (E) gCAMP6s expression in VIP interneurons. (F) Ca^2+^ transients in VIP interneurons at L-on. (G) Left: Histogram of modulation index for VIP cells around L-on. Controls: 223 cells, 6 mice. Mutants: 87 cells, 3 mice. Right: Average modulation index for VIP interneurons around L-on. Error bars and shading show s.e.m., p**<0.01, p***<0.001.

To examine how changes in VIP-INs activity might contribute to loss of state-dependent cortical modulation, we compared VIP interneuron activation patterns in mutants and controls. We expressed the genetic calcium indicator gCaMP6s in VIP-INs and used 2-photon imaging (Figures 3E, S6) to assay their activity during behavioral state transitions. VIP-INs increased their activity around locomotion onset in control mice, but their state-dependent modulation was reduced in mutants (Figures 3F-G, S6D). The recruitment of VIP-IN activity by excitatory afferents may thus be compromised. Whole-cell recordings from acute brain slices confirmed reduced glutamatergic input to VIP-INs in the mutants (Figure S6A-B).

We next compared the state-dependent effects of VIP-IN disruption on putative excitatory and inhibitory neurons. In contrast to the complete loss of modulation observed in RS cells, the state-dependence of FS cell firing was largely unchanged in mutants as compared to controls (Figure 4). FS cells in both mutants and controls showed robust increases in firing rate at locomotion onset (Figure 4A,B,D) and brief suppression of firing rate at locomotion offset relative to later quiescent periods (Figure 4C-E). These data further support the idea that developmental VIP-IN disruption decouples excitatory and inhibitory cortical networks.

**Figure 4.**
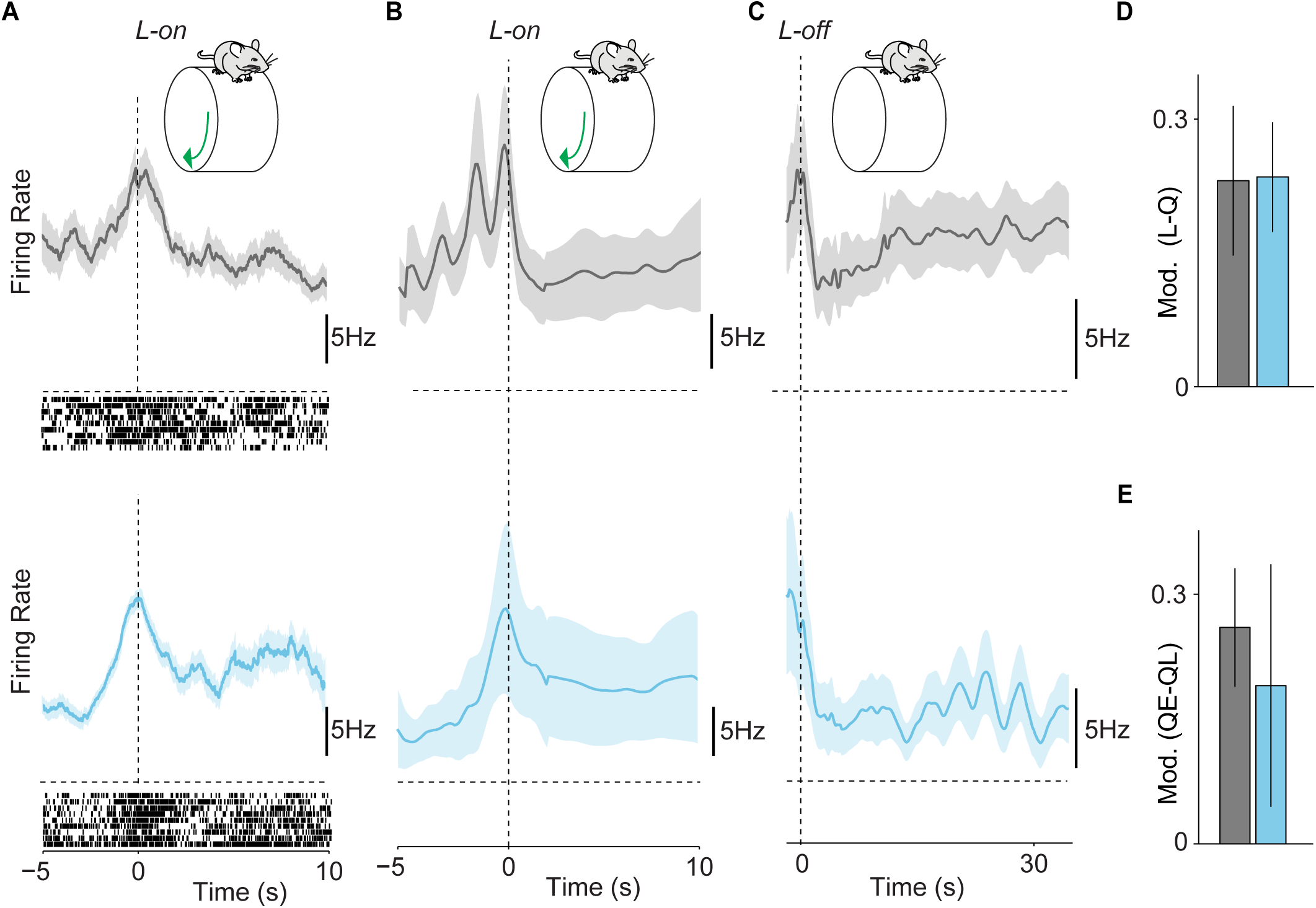
State modulation of FS cell activity in ErbB4-VIP mutants is unaffected. (A) Average firing rate for example FS cells around locomotion onset (L-on) in controls and mutants. (B) Population average change in firing rate around locomotion onset for FS cells in controls and mutants. (C) (B) Population average change in firing rate around locomotion offset for FS cells in controls and mutants. (D) Firing rate modulation in early locomotion (L; -0.5 to 0.5s around L-on) period as compared to quiescence (Q). (E) Firing rate modulation in late (QL) vs. early (QE) quiescence. Controls: 21 cells, 8 animals. Mutants: 29 cells, 6 animals. There were no significant differences between control and mutant mice.

### Developmental disruption of VIP interneurons compromises sensory processing and learning

Schizophrenic patients exhibit deficits in basic visual processing and perception, including reduced contrast sensitivity (Slaghuis, 2004; Martinez et al., 2008; Tan et al., 2013; Serrano-Pedraza et al., 2014), functions that rely on V1. To test the impact of ErbB4 deletion on cortical visual processing, we performed extracellular recordings in V1 of awake behaving animals during visual stimulation. In mutants, visual responses to drifting grating stimuli were reduced in amplitude in RS, but not FS cells (Figures 5A-D, S7A). Neither the observed decrease in state modulation nor the reduced visual responsiveness was correlated with elevated firing rates on a cell-by-cell basis (Figure S8).

**Figure 5.**
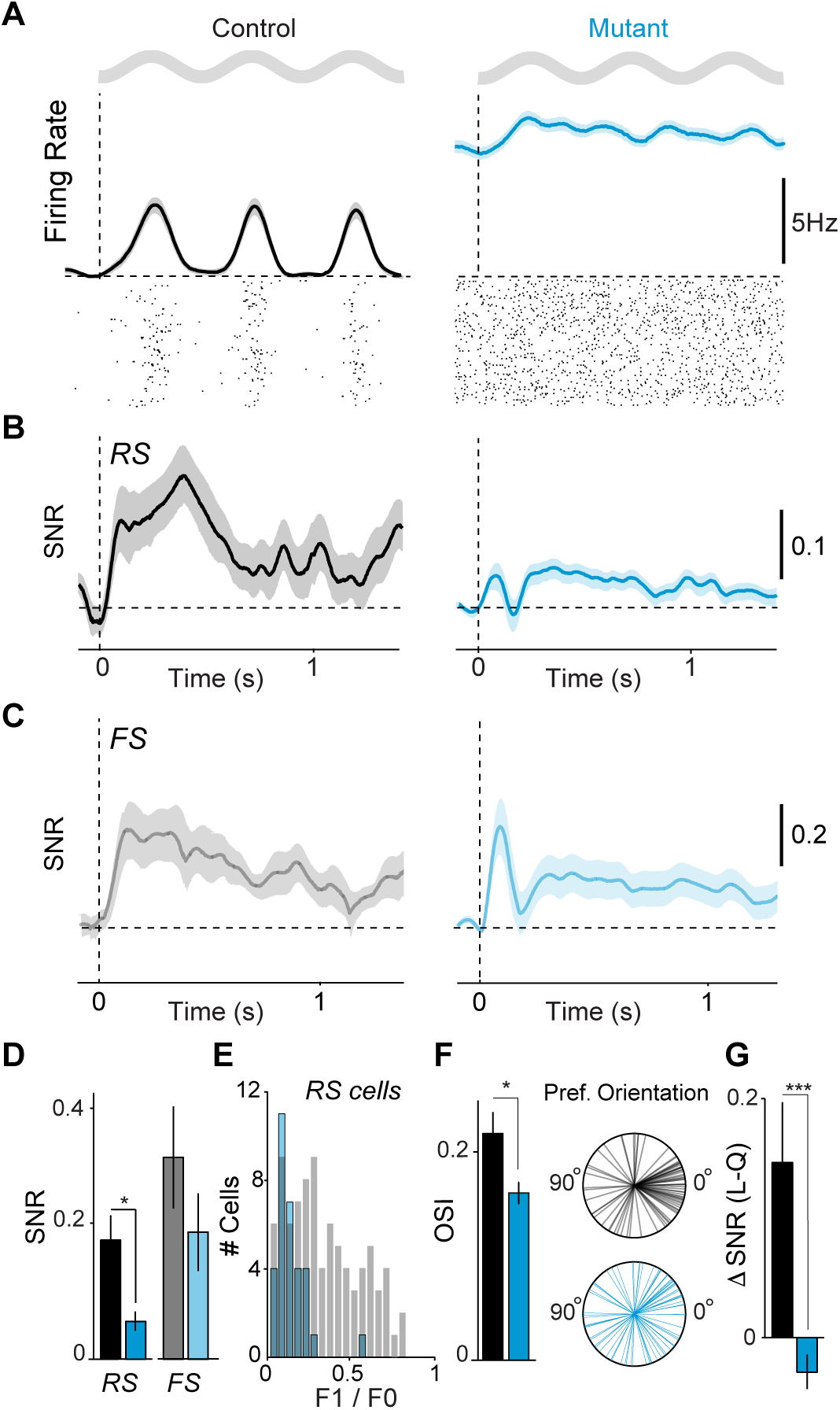
ErbB4 mutants exhibit reduced visual response selectivity. (A) Average firing rate and spike raster plot for example RS cells in response to a drifting grating stimulus in controls and mutants. (B) Average rate modulation relative to inter-trial interval around stimulus onset for RS cells. Controls: 106 RS, 8 mice. Mutants: 92 RS, 7 mice. (C) Average rate modulation relative to inter-trial interval around stimulus onset for RS cells. Controls: 21 cells, 8 animals. Mutants: 28 cells, 7 animals. (D) Signal-to-noise ratio of visual responses for RS and FS cells. (E) Histogram of F1/F0 values for RS cells. (F) Average orientation selectivity index (OSI; left) of all RS cells. Radial plots of preferred orientations of all RS cells (right). Controls: 55 cells, 7 mice. Mutants: 61 cells, 8 mice. Panels A-F measured during quiescence. (G) Increase in stimulus rate modulation of RS cells during locomotion as compared to quiescence.

In addition to changes in overall visual responsiveness, receptive field properties of V1 neurons were altered. RS cells in mutants exhibited less linear summation properties than controls, leading to an overall population shift towards low F1/F0 values associated with complex, rather than simple, cells (Figure 5E, S7F). RS cells in mutants were less orientation selective than those in controls (Figure 5F, S7C). Furthermore, whereas neurons in control mice showed a significant bias towards horizontal (0° angle) stimuli, a feature of cortical visual tuning that is refined after eye opening (Rochefort et al., 2011), this bias was absent in mutant mice (Figures 5F, S7D, E). We found that deletion of ErbB4 from VIP interneurons also eliminated the increase in visual response gain normally observed during periods of locomotion and arousal. We and others have previously shown that the signal-to-noise of visual responses in controls increases in the active state (Niell and Stryker, 2010; Fu et al., 2014; Vinck et al., 2015), but this enhancement was absent in the mutants (Figure 5G). Because visual response properties emerge from the establishment of appropriate synaptic connections, these findings suggest that deletion of ErbB4 from VIP-INs has a broad impact on the development of cortical synaptic interactions.

To directly test whether the altered spontaneous and visually evoked cortical activity observed in mutants contributed to perceptual deficits, we trained mice to perform a visual detection task in which stimulus contrast varied (Figure 6A). Mutants and controls had similar detection rates for high-contrast stimuli and similar false alarm rates (Figure 6B-C). The two groups performed equally well during early training sessions and showed equivalent initial psychometric functions (Figure 6D-E, S9D). However, control mice showed improved performance for low-contrast stimuli over several training days but mutant mice did not (Figure 6B,D-E, S9E), suggesting that developmental dysregulation of VIP-INs disrupts both neural activity and sensory perceptual learning, consistent with deficits seen in schizophrenia.

**Figure 6.**
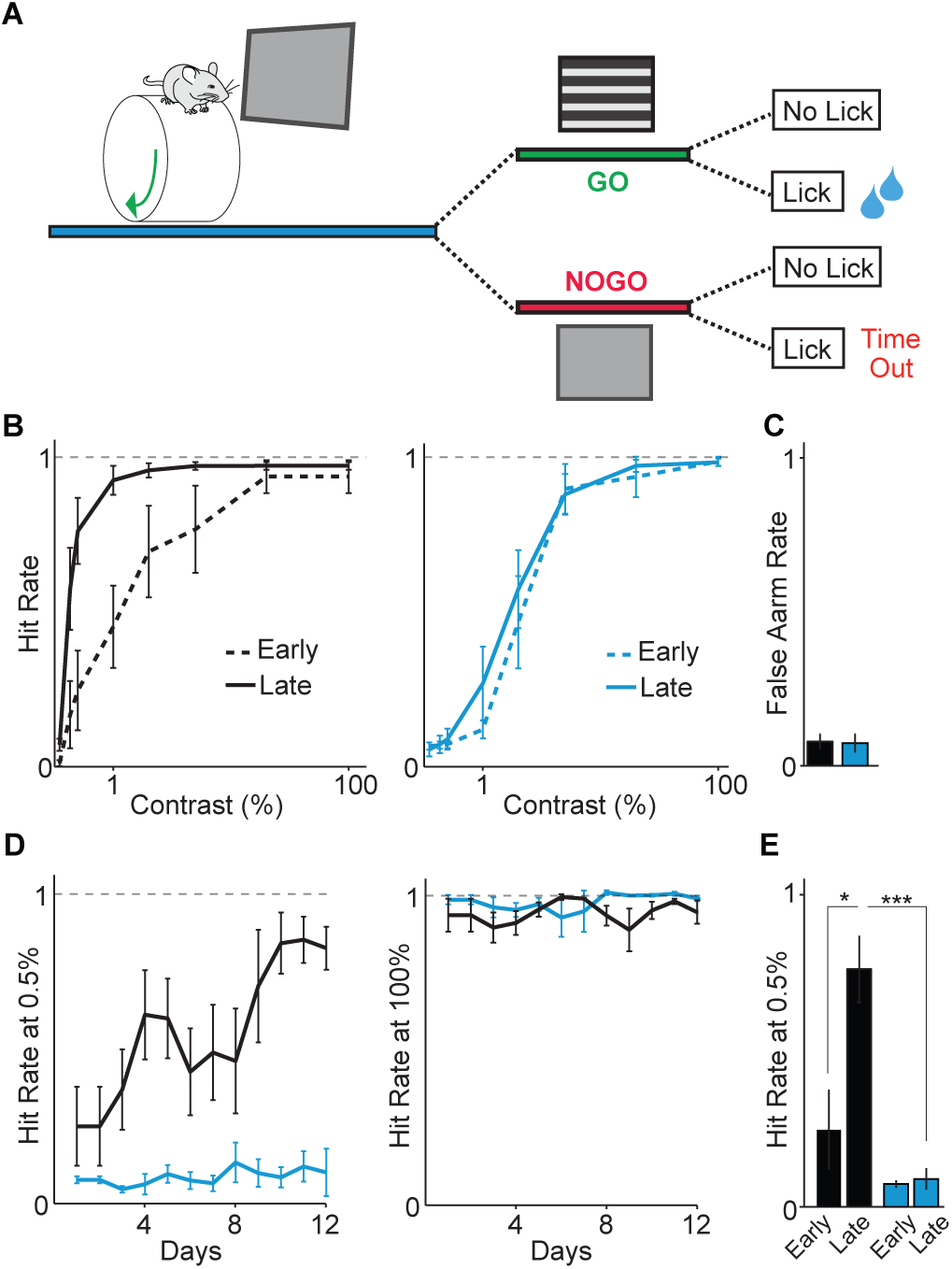
VIP interneuron disruption causes impaired sensory learning. (A) Schematic of visual detection task. On each trial, either a grating appeared (GO) or there was no change (NOGO). Correct hits were rewarded with water, whereas incorrect hits were followed by a time-out. Controls: 4 mice. Mutants: 4 mice. (B) Control and mutant psychophysical performance curves for early and late training sessions. (C) Average false alarm rates. (D) Average performance for low and high contrast stimuli.(E) Average performance at low contrast for early and late training days. All panels: error bars and shading show s.e.m.; p*<0.05, p***<0.001.

## DISCUSSION

Schizophrenia is associated with increased cortical and hippocampal excitability, reduced gamma oscillations and neural synchrony, and altered sensory responses. Current evidence suggests that these deficits may be the result of disrupted GABAergic inhibition, which normally acts in balanced coordination with glutamatergic excitation. Synthesis and reuptake of GABA and total numbers of both inhibitory interneurons and inhibitory synapses are reduced in schizophrenic patients (Lewis et al., 1999; Pierri et al., 1999; Lewis et al., 2004; Lewis et al., 2005; Konopaske et al., 2006; Hashimoto et al., 2008; Cruz et al., 2009), suggesting overall dysregulation of inhibition. These changes disrupt cortical circuits underlying cognition and perception, functions that are widely compromised in schizophrenic patients. Although most previous work has focused on the role of PV interneuron disruption in disease-related deficits, the diversity of GABAergic interneurons and their functional roles in neural circuits suggests that other inhibitory cell types, such as VIP interneurons, may be important in the pathophysiology of this complex disease.

We used deletion of the schizophrenia risk gene ErbB4 to test the role of VIP-INs in disease-related neural and behavioral deficits. Although VIP-INs represent only ~12% of cortical GABAergic cells (Rudy et al., 2011), their developmental dysregulation had a surprising, long-term impact on cortical function. ErbB4 deletion from VIP-INs caused increased excitability of putative excitatory cortical neurons, decreased synchrony and participation in gamma rhythms, and reduced cortical sensory responses. These neural changes were associated with deficits in performance of a visual perception task and other behavioral assays. Disruption of the Nrg1-ErbB4 signaling pathway in VIP-INs may thus contribute to key elements of neurodevelopmental pathophysiology.

Recent work has suggested a model where VIP-INs regulate cortical circuits predominantly by inhibiting other interneurons, including the dendrite-targeting SOM cells (Pfeffer et al., 2013), thereby disinhibiting pyramidal neurons (though see Polack et al., 2013 and Dipoppa et al., 2016 for a different conclusion). We found that VIP-INs in mutants received decreased excitatory inputs and were not appropriately activated at locomotion onset, suggesting that their impact on the local neural circuit was diminished. However, this decrease in VIP activation was associated with a sustained increase, rather than a decrease, in firing of putative pyramidal neurons. In contrast, the activity of FS interneurons was unaffected. These results suggest that imbalances of excitation and inhibition in cortical circuits may result from elevated excitatory neuron activity, independent of changes in fast inhibition.

Fast-spiking interneurons play a key role in regulating the temporal pattern of excitatory cortical activity, and contribute to the generation of gamma oscillations and spike synchrony (Cardin et al., 2009; Sohal et al., 2009). In healthy cortical circuits, FS interneurons entrain excitatory neurons to the gamma rhythm and their activity is tightly coupled to that of excitatory neurons and other FS cells (Hasenstaub et al., 2005; Cardin et al., 2009; Vinck et al., 2013). These fine time-scale interactions are critical for restricting the generation of excitatory action potentials (Pouille and Scanziani, 2001; Wehr and Zador, 2003; Cardin et al., 2010) and for information encoding and transmission across long-range circuits (Fries, 2009). Surprisingly, following deletion of ErbB4 from VIP-INs, RS and FS populations were decoupled. Our findings suggest that VIP-INs play an unanticipated and uniquely influential role in cortical circuits, and developmental dysregulation of the Nrg-1/ErbB4 signaling pathway in these cells permanently compromises the ability of cortical circuits to participate in the encoding and transmission of sensory information.

Cortical activity patterns and sensory responses are strongly modulated by behavioral state, such as sleep, wakefulness, and attention. This modulation is thought to be important for enhancing encoding of behaviorally relevant information and is compromised in schizophrenia (Lewis and Lieberman, 2000). Previous work has highlighted a role for VIP-INs in the state-dependent modulation of cortical gain control (Fu et al., 2014). ErbB4 deletions from VIP-INs caused a loss of the enhanced visual response gain normally associated with arousal and locomotion (Saleem et al., 2013; Erisken et al., 2014; Vinck et al., 2015; Mineault et al., 2016) suggesting that developmental VIP-INs disruption decreases the dynamic range of cortical circuit activity. However, disruption of VIP-INs did not reduce the transmission of behavioral state-related signals to fast-spiking interneurons. Developmental impairment of VIP-INs thus interrupts the ability of cortical circuits to adapt rapidly to ongoing cognitive and behavioral demands and to encode information about behaviorally relevant environmental features.

Previous work has found that global disruption of Nrg-1 or ErbB4 impairs normal synaptic function and plasticity (Kwon et al., 2005; Hahn et al., 2006; Chen et al., 2010; Abe et al., 2011; Geddes et al., 2011; Pitcher et al., 2011; Shamir et al., 2012) and alters the development of excitatory synapses onto PV interneurons and PV synapses onto pyramidal neurons (Li et al., 2007; Fazzari et al., 2010). GABAergic inhibition plays a key role in the early postnatal development of appropriate synaptic receptive field structure (Hensch, 2005) and in the induction of plasticity in mature circuits (van Versendaal and Levelt, 2016), including via VIP-INs (Fu et al., 2015; Mardinly et al., 2016). VIP-specific ErbB4 deletion decreased visual responsiveness and altered the response properties of putative excitatory neurons, suggesting that developmental dysregulation of VIP-INs disrupts the establishment or maintenance of receptive field properties in visual cortex neurons.

The disruption of network activity and sensory processing we observed following ErbB4 deletion was associated with compromised visual task performance and a consistently elevated threshold for visual contrast detection. It is unlikely that the deficits observed in the mutants resulted from failure to see stimuli or perform a motor response, as the two groups demonstrated similar baseline psychophysical performance. In comparison, deletion of ErbB4 from SST-expressing GABAergic neurons in the thalamic reticular nucleus impairs attentional switching but not learning or performance of a basic sensory detection task (Ahrens et al., 2015). Impairments following ErbB4 deletion from VIP-INs were not restricted to visual functions. Mutants showed locomotor hyperactivity and a deficit in marble burying, two behavioral phenotypes observed previously in ErbB4 deletion mice and other genetic models of schizophrenia (Leung and Jia, 2016). Together, these results suggest that behavioral and perceptual deficits following global disruption of Nrg-1/ErbB4 signaling may not result solely from PV interneuron disruption, but instead arise from the summed impact of multiple GABAergic mechanisms.

In summary, we find a potential role for VIP interneurons in the pathophysiology underlying neurodevelopmental dysregulation in schizophrenia. This contrasts with the prevailing view, which focuses almost exclusively on a role for PV interneurons (Rico and Marin, 2011; Del Pino et al., 2013). The severe disruption of cortical function following selective early mutation of VIP interneurons further suggests an unanticipated role for these cells in normal cortical development and refinement during learning. Despite being few in number, VIP interneurons are targets for multiple neuromodulatory systems and potent regulators of cortical activity. VIP interneuron disruption may thus be a powerful mechanism underlying the large-scale dysregulations of cortical activity patterns, behavioral and sensory modulation, and perceptual function that are hallmarks of psychiatric disease.

## Author Contributions

RBB, MV, KAF, and JAC designed experiments. RBB and MV performed *in vivo* electrophysiology recordings. RBB and MV performed visual behavior experiments. RBB and MV analyzed *in vivo* electrophysiology and visual behavior data. GL and KAF performed surgical implants for 2-photon imaging. KAF performed 2--photon imaging experiments and analyzed data. DL and MJH performed and analyzed *in vitro* electrophysiology experiments. RBB, MV, and JAC wrote manuscript.

## Acknowledgements

The authors are grateful to A. Buonanno for the ErbB4 antibody and A. Koleske for the ErbB4^F/F^ mice. We thank R. Pant for assistance with histology and A. Airhart, T. Church, V. Hernandez, J. Mossner, and A. Murray for assistance with behavioral assays. We thank U. Knoblich for initial work on the behavioral apparatus. We thank Q. Perrenoud for mouse illustrations. This work was supported by a Brown-Coxe fellowship, a Jane Coffin Childs Fellowship, and a NARSAD Young Investigator Award to RBB; a Rubicon fellowship and a Human Frontiers Postdoctoral Fellowship to MV; NIH R01 MH099045 to MJH; and NIH R01 MH102365, a Smith Family Award for Excellence in Biomedical Research, a Klingenstein Fellowship Award, an Alfred P. Sloan Fellowship, a NARSAD Young Investigator Award, and a McKnight Fellowship to JAC.

## EXPERIMENTAL PROCEDURES

### Mouse Strains

All animal handling and maintenance was performed according to the regulations of the Institutional Animal Care and Use Committee of the Yale University School of Medicine. *ErbB4^F/F^* mice were crossed to *ErbB4^F/+^ VIP^Cre^* to generate mutant animals (*ErbB4^F/F^ VIP^Cre^*) and littermate controls (*ErbB4^F/F^*, or *VIP*^*Cre*^). Crosses for histology included the *Ai9* reporter line.

### Immunohistochemistry

Control mice *Dlx6a*^*Cre*^ Ai9, *PV*^*Cre*^ Ai9, *SST*^*Cre*^ Ai9, or *VIP*^*Cre*^ Ai9, and mutant *ErbB4^F/F^ VIP^Cre^* Ai9 animals were examined using immunohistochemistry. Brains were fixed by transcardial perfusion with 4% paraformaldehyde (PFA)/phosphate buffered saline (PBS) solution followed by a 1 hour post-fixation on ice with 4% PFA/PBS solution. Brains were rinsed with PBS and cryoprotected by using 15% sucrose/PBS solution for 6 hours and 30% sucrose/PBS solution overnight at 4°C. Tissues were embedded in Tissue Tek, frozen on dry ice, and cryosectioned at 20 µm thickness. Sections for immunohistochemistry analysis were processed using 1.5% normal goat serum (NGS) and 0.1% Triton X-100 in all procedures except washing steps, where only PBS was used. Sections were blocked for 1 hour, followed by incubation with the primary antibodies overnight at 4°C. Cryostat tissue sections were stained with the primary antibody rabbit anti-ErbB4 (1:2000; mAB10 courtesy of A. Buonanno). Secondary antibody conjugated with Alexa fluorescent dyes 488 (Invitrogen) raised from goat was applied for 1 hr at room temperature for visualizing the signals. Nuclear counterstaining was performed with DAPI solution.

All analysis was evaluated in the primary visual cortex (V1). To minimize counting bias we compared sections of equivalent bregma positions, defined according to the Mouse Brain atlas (Paxinos and Franklin, 2001). The total number of cells expressing tdTomato (from the *Ai9* reporter mouse line) were counted for a defined optical area. The percentages of cortical interneurons expressing ERBB4 or subtype specific markers (DLX6, PV, SST, VIP) among fate-mapped cells were calculated as a ratio between the number of double positive cells (Marker and tdTomato) over the total number of tdTomato positive cells. All data were represented as mean ± SEM, unpaired Student’s t-test.

### Headpost surgery and wheel training

Mice were handled for 5-10 min/day for 5 days prior to the headpost surgery. On the day of the surgery, the mouse was anesthetized with isoflurane and the scalp was shaved and cleaned three times with Betadine solution. An incision was made at the midline and the scalp resected to each side to leave an open area of skull. Two skull screws (McMaster-Carr) were placed at the anterior and posterior poles. Two nuts (McMaster-Carr) were glued in place over the bregma point with cyanoacrylate and secured with C&B-Metabond (Butler Schein). The Metabond was extended along the sides and back of the skull to cover each screw, leaving a bilateral window of skull uncovered over primary visual cortex. The exposed skull was covered by a layer of cyanoacrylate. The skin was then glued to the edge of the Metabond with cyanoacrylate. Analgesics were given immediately after the surgery and on the two following days to aid recovery. Mice were given a course of antibiotics (Sulfatrim, Butler Schein) to prevent infection and were allowed to recover for 3-5 days following implant surgery before beginning wheel training.

Once recovered from the surgery, mice were trained with a headpost on the wheel apparatus. The mouse wheel apparatus was 3D-printed (Shapeways Inc.) in plastic with a 15 cm diameter and integrated axle and was spring-mounted on a fixed base. A programmable magnetic angle sensor (Digikey) was attached for continuous monitoring of wheel motion. Headposts were custom-designed to mimic the natural head angle of the running mouse, and mice were mounted with the center of the body at the apex of the wheel. On each training day, a headpost was attached to the implanted nuts with two screws (McMaster-Carr). The headpost was then secured with thumb screws at two points on the wheel. Mice were headposted in place for increasing intervals on each successive day. If signs of anxiety or distress were noted, the mouse was removed from the headpost and the training interval was not lengthened on the next day. Mice were trained on the wheel for up to 7 days or until they exhibited robust bouts of running activity during each session. Mice that continued to exhibit signs of distress were not used for awake electrophysiology sessions.

#### *In vivo* electrophysiology

All extracellular single-unit, multi-unit, and LFP recordings were made with an array of independently moveable tetrodes mounted in an Eckhorn Microdrive (Thomas Recording). Signals were digitized and recorded by a Digital Lynx system (Neuralynx). All data were sampled at 40kHz. All LFP recordings were referenced to the surface of the cortex. LFP data were recorded with open filters and single unit data was filtered from 600-9000Hz.

Awake recordings were made from mice that had received handling and wheel training as described above. On the initial recording day, a small craniotomy was made over V1 under light isoflurane anesthesia. The craniotomy was then covered with Kwik-Cast (World Precision Instruments), after which the mouse was allowed to recover for 2 hours. Mice were then fitted with a headpost and secured in place on the wheel apparatus before electrodes were lowered. Recording electrodes were initially lowered to ~150 µm, then independently adjusted after a recovery period of 30-60 min. At the end of a recording session, the craniotomy was flushed with saline and capped. On subsequent recording days, the craniotomy was flushed with saline before placing the electrode array in a new site. Recordings were performed mainly in the second half of the light portion of the light/dark cycle.

#### *In vitro* electrophysiology

Under isoflurane anesthesia, mice were decapitated and transcardially perfused with ice-cold choline-artificial cerebrospinal fluid (choline-ACSF) containing (in mM): 110 choline, 25 NaHCO_3_, 1.25 NaH_2_PO_4_, 2.5 KCl, 7 MgCl_2_, 0.5 CaCl_2_, 20 glucose, 11.6 sodium ascorbate, 3.1 sodium pyruvate. Acute occipital slices (300 µm) were prepared from the left hemisphere and transferred to ACSF solution containing (in mM): 127 NaCl, 25 NaHCO_3_, 1.25 NaH_2_PO_4_, 2.5 KCl, 1 MgCl_2_, 2 CaCl_2_, and 20 glucose bubbled with 95% O_2_ and 5% CO_2_. After an incubation period of 30 min at 32˚C, the slices were maintained at room temperature until use.

Visualized whole-cell recordings were performed by targeting fluorescently labeled VIP-INs in the monocular region of the primary visual cortex (V1). All recordings were performed at room temperature. Criteria for recording included a series resistance (Rs) of <20 MΩ. For miniature excitatory postsynaptic current recordings, the ACSF contained 1 µM TTX to block sodium channels. The internal solution contained (in mM): 126 cesium gluconate, 10 HEPES, 10 sodium phosphocreatine, 4 MgCl_2_, 4 Na_2_ATP, 0.4 Na_2_GTP, 1 EGTA (pH 7.3 with CsOH). Cells were voltage-clamped at -70 mV.

### *In vivo* imaging

To express the genetically encoded calcium indicator GCaMP6s, mice were anesthetized with 1-2% isoflurane mixed with pure oxygen and a small craniotomy was made over primary visual cortex. Each mouse received two 100 nl injections of adenoassociated virus (AAV2.5-synapsin1-GCaMP6s, University of Pennsylvania Vector Core) at coordinates (in mm from Bregma): AP -3.5, ML 1.8, DV 0.45 and AP -2.8, ML 2.5, DV 0.45. Injections were made via beveled glass micropipette at a rate of ~10 nl/min. After injection, pipettes were left in the brain for ~5 minutes to prevent backflow. Imaging experiments were conducted 25-30 days after virus injection. For implantation of the imaging window, mice were anesthetized using a mixture of ketamine (80 mg/kg) and xylazine (5 mg/kg), and a ~2 mm diameter craniotomy was opened over V1. An imaging window consisting of a small rectangular glass piece attached to a 5mm circular cover glass using an ultraviolet-curing adhesive (Norland Products) was inserted into the craniotomy and secured to the skull with Metabond. A custom titanium head post was secured to the skull with Metabond.

Imaging was performed using a resonant scanner-based two-photon microscope (MOM, Sutter Instruments) coupled to a Ti:Sapphire laser (MaiTai DeepSee, Spectra Physics) tuned to 940 nm for GCaMP6. Emitted light was collected using a 25x 1.05 NA objective (Olympus). Mice were placed on the wheel and head-fixed under the microscope objective. To prevent light contamination from the display monitor, the microscope was enclosed in blackout material that extended to the headpost. Images were acquired using ScanImage 4.2 at ~30 Hz, 256x256 pixels (290x290 um). Imaging of layer 2/3 was performed at ~150-300 um depth relative to the brain surface. Images were continuously monitored throughout the experiments, and slow drifts of the image were manually corrected. For each mouse, 1-4 fields of view were imaged.

### Visual stimulation

Visual stimuli were presented on an LCD monitor at a spatial resolution of 1680x1050, a real-time frame rate of 60Hz, and a mean luminance of 45 cd/m^2^ positioned 15cm from the eye. The LCD monitor used for visual stimulation (22 inches) was mounted on an arm and positioned on the right side of the animal, perpendicular to the surface of the right eye. The screen was placed so that stimuli were only presented to the right eye. Stimuli were generated by custom-written software (J. Cardin, Matlab). Initial hand-mapping was performed to localize the receptive fields of identified cells. To maximize data collection, visual stimuli were positioned to cover as many identified receptive fields as possible. All stimuli were sinusoidal drifting gratings at a temporal frequency of 2 Hz, presented at a fixed duration of 1.5 s with an interstimulus interval of 2 s. We used blocks of visual stimuli where contrast was held at 100% and orientation was varied. To determine orientation tuning, gratings were presented at 12 different orientations, randomized and presented 20-50 times per orientation. Orientation tuned stimuli were optimized for mean spatial frequency.

### Data analysis

Spikes were clustered semi-automatically using the following procedure. We first used the KlustaKwik 2.0 software to identify a maximum of 30 clusters using the waveform Energy and Energy of the waveform’s first derivative as clustering features. We then used a modified version of the M-Clust environment to manually separate units and we selected well isolated units. We further ensured that maximum contamination of the ISI (Inter-spike-interval) histogram <1.5 ms was smaller than 0.1%. In a small number of cases we accepted clusters with isolation distances smaller than 20, which could be caused by e.g. non-Gaussian clusters, only if separation was of sufficient quality as judged by comparing the cluster with all other noise clusters (20%, 80% quantiles of ID = 19, 43; median±standard error of median = 25±0.5). Unit data were analyzed for firing rates and patterns, correlations, and visual responses using custom-written Matlab software (see Supplemental Information).

### Visual detection task

Mice were trained to perform a GO-NOGO visual contrast detection task while head-fixed on a wheel. The screen was placed ~15cm from the mouse. During initial shaping stages, mice were trained to lick a water spout in response to presentation of a high-contrast, full-screen stimulus. A tone cue was given to signal the onset of each trial. When a performance criterion of >80% hit rates and <20% false alarm rates was reached, they were moved to the full version of the task where the stimulus contrast varied randomly across trials. Stimulus contrast was selected on each trial from the series: 0, 0.35, 0.425, 0.5, 1, 2, 5, 20, 100%. A correct response (hit) was rewarded with a small (~2µl) drop of water. False alarms were punished with a bright screen, a high-frequency tone, and an extended inter-trial interval between 15 and 17s. Correct and incorrect rejections (miss) were neither rewarded nor punished but were followed by an inter-trial-interval between 5 and 7s. Mice were put on the task for a total of 45 minutes per session.

---

**Figure S1 (Related to Figure 1).**
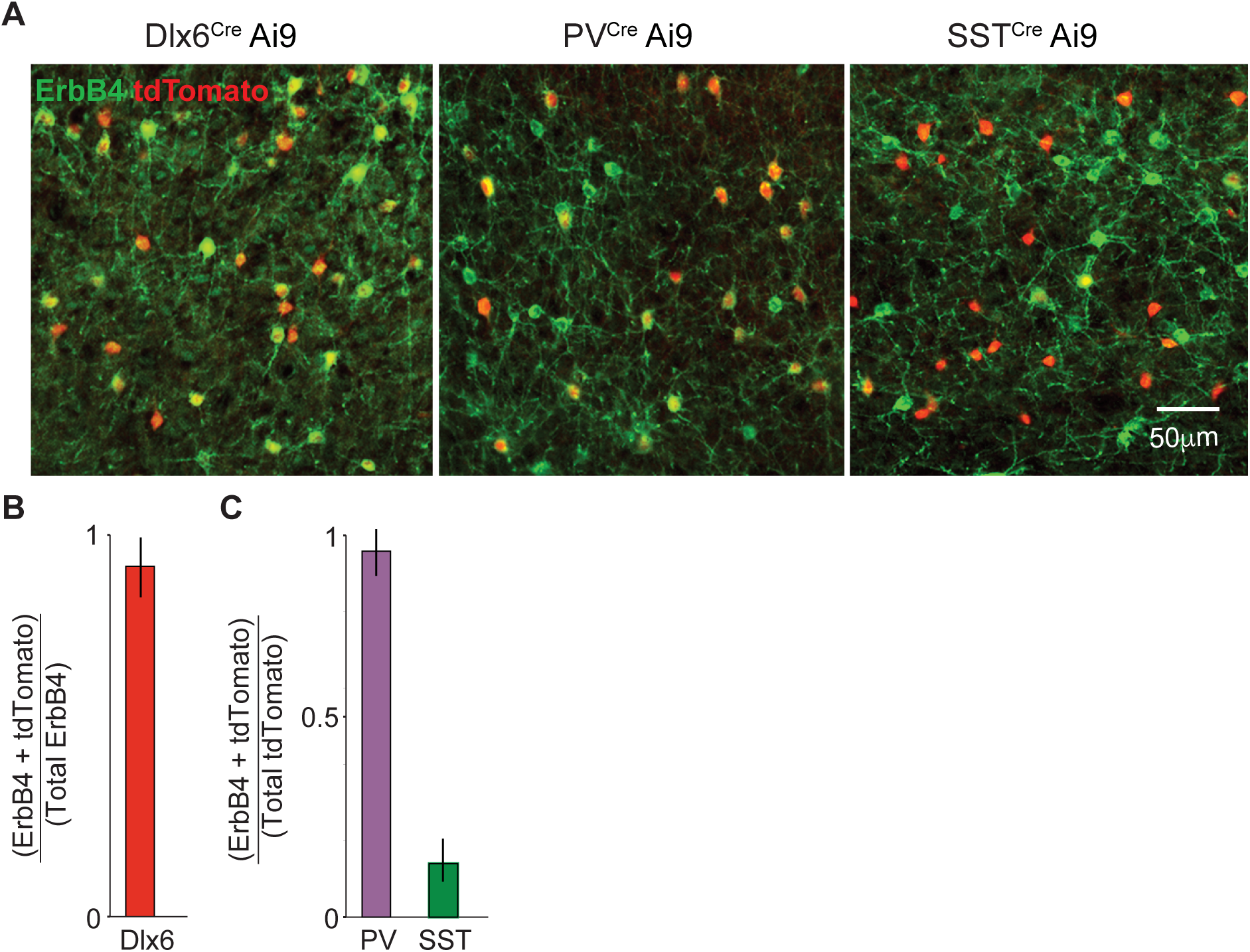
Expression of ErbB4 in interneuron populations. (**A**) ErbB4 (green) and tdTomato (red) immunohistochemistry in the visual cortex (V1) of P60 Dlx6^Cre^ (pan-interneuron marker), PV^Cre^ and SST^Cre^ mice crossed to the reporter line Ai9. (**B**) ErbB4 expression in Dlx6 fate-mapped cells shows that the vast majority of ErbB4 expressing cells in the V1 are interneurons. (**C**) PV and SST fate-mapped interneurons that express ErbB4 in the V1. Most PV interneurons, but few SST interneurons, express ErbB4. N=6 mice for each of the quantifications. Error bars indicate mean ± s.e.m.

**Figure S2 (Related to Figure 1).**
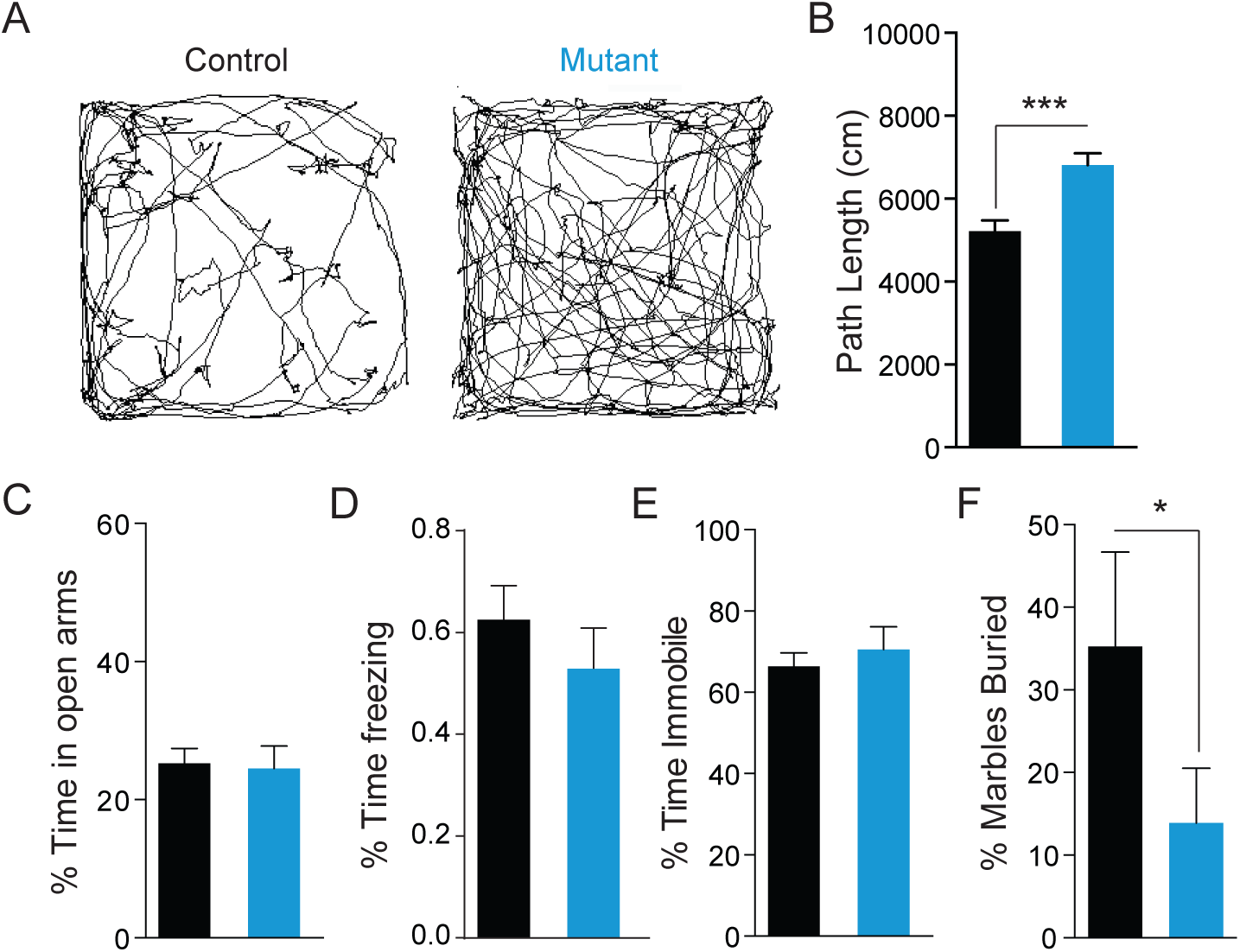
Behavioral characterization of ErbB4-VIP mutants and controls. (**A**) Example paths in the first five minutes of the open field assay for a control (left) and mutant (right). (**B**) Population quantification of total open field path length (n = 29 controls, 18 mutants). (**C**) Time spent in the open arms of the elevated plus maze assay (n= 29 controls, 21 mutants). (**D**) Percentage of time freezing 24h after the contextual fear conditioning paradigm (n = 11 controls, 15 mutants). (**E**) Percentage of time spent immobile in the forced swim assay (n = 16 controls, 6 mutants). (**F**) Percentage of marbles buried in the marble burying assay (n = 14 controls, 6 mutants). Error bars indicate mean ± s.e.m. P * < 0.05, ***<0.001.

**Figure S3 (Related to Figure 1).**
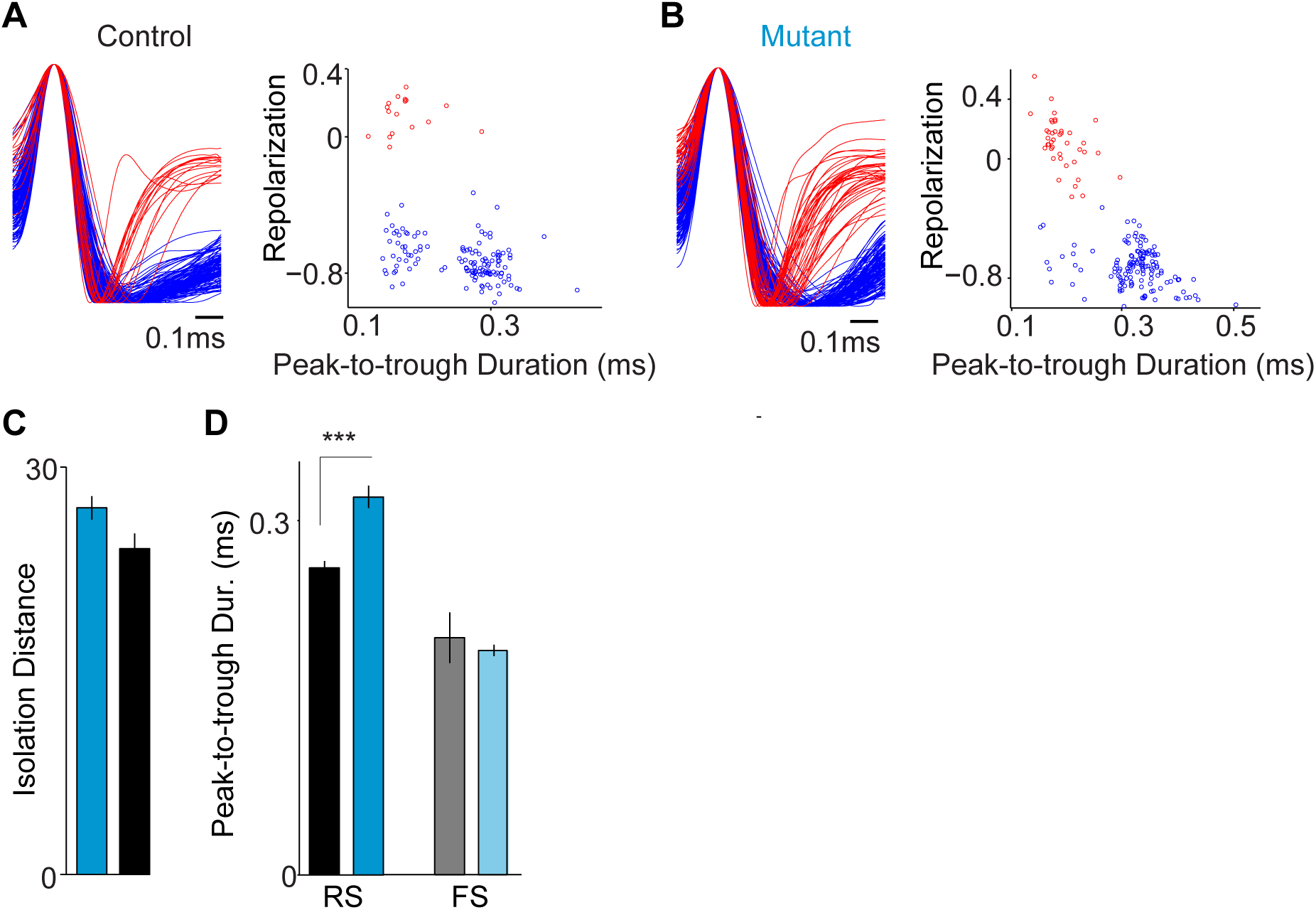
Waveform separation and characteristics for ErbB4-VIP mutants and controls. (**A**) Left: normalized AP waveforms, aligned to AP peak. Right: peak-to-trough duration of action potentials (APs) vs. AP repolarization metric, which was defined as the value of the normalized (between -1 and +1) AP waveform at 450 ms (as in Vinck et al., 2015). We separated three clusters of waveforms in both controls and mutants using a Gaussian mixture model, as in Vinck et al. (2015). Putative RS cells can be subdivided into groups with thin and broad spikes (Haider et al., 2010, Vinck et al., 2015) that both have low firing rates in wild type animals (Vinck et al., 2015). The distributions of RS and FS cell firing rates were largely non-overlapping in control mice (see Fig. 1). (**B**) Same as in (**A**), but for mutant mice. (**C**) Median ± s.e.m. of isolation distance for cells in mutant and control mice. (**D**) Peak-to-trough duration for RS and FS cells. RS cells in mutant mice have broader action potentials on average. Controls: 153 RS, 15 FS cells, 8 mice. Mutants: 134 RS, 32 FS cells, 8 mice. Error bars indicate mean ± s.e.m. P***<0.001.

**Figure S4 (Related to Figure 1).**
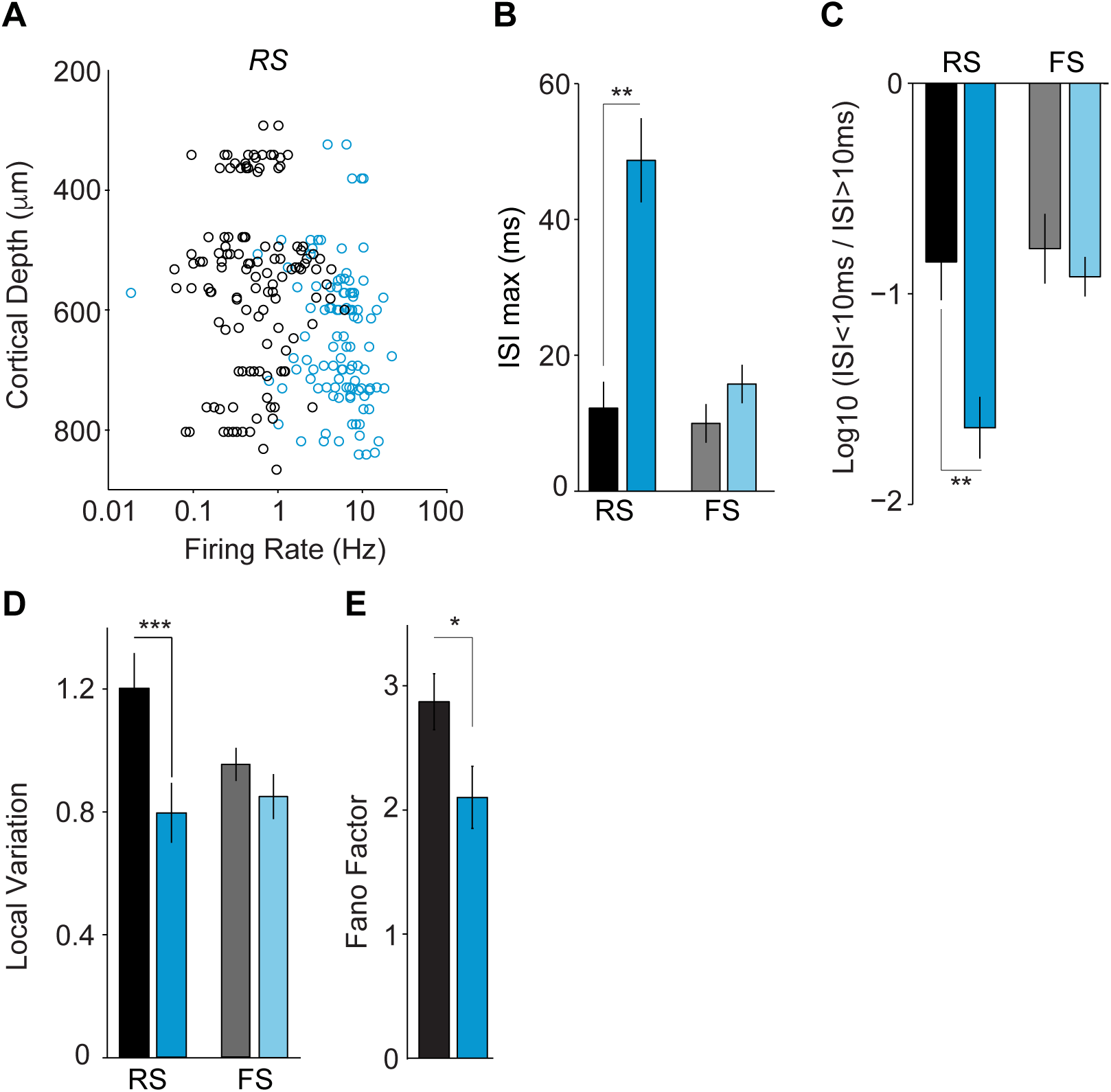
Additional measures of the impact of ErbB4-VIP deletion further support a cell type-specific impact on RS firing. (**A**) Firing rate (Hz) vs cortical depth for RS cells in control (black) and mutant (cyan) mice. (**B**) Peak of inter-spike interval (ISI) histogram for RS and FS cells during quiescence in controls (black) and mutants (cyan). We determined the peak time of the ISI histogram separately for each cell based on an ISI histogram that extended up to 1s in 1ms bins. RS, but not FS cells, have a later peak ISI peak time in mutants. (**C**) Log ratio of number of ISIs <10ms over the number of ISIs >10ms, a measure of burstiness. Higher values indicate more burstiness. RS, but not FS, cells in control animals have a larger percentage of short ISIs as compared to mutants. (**D**) Local Variation, which is a measure of the irregularity of spike trains. RS, but not FS cells in mutant animals fire more regularly than those in controls. (**E**) Average Fano factor, defined as the variance of the spike count over the mean spike count, for RS cells. Controls: 153 RS, 15 FS cells, 8 mice. Mutants: 134 RS, 32 FS cells, 8 mice. (**B-E**) Error bars indicate mean ± s.e.m. p**<0.01, p***<0.001.

**Figure S5 (Related to Figure 2).**
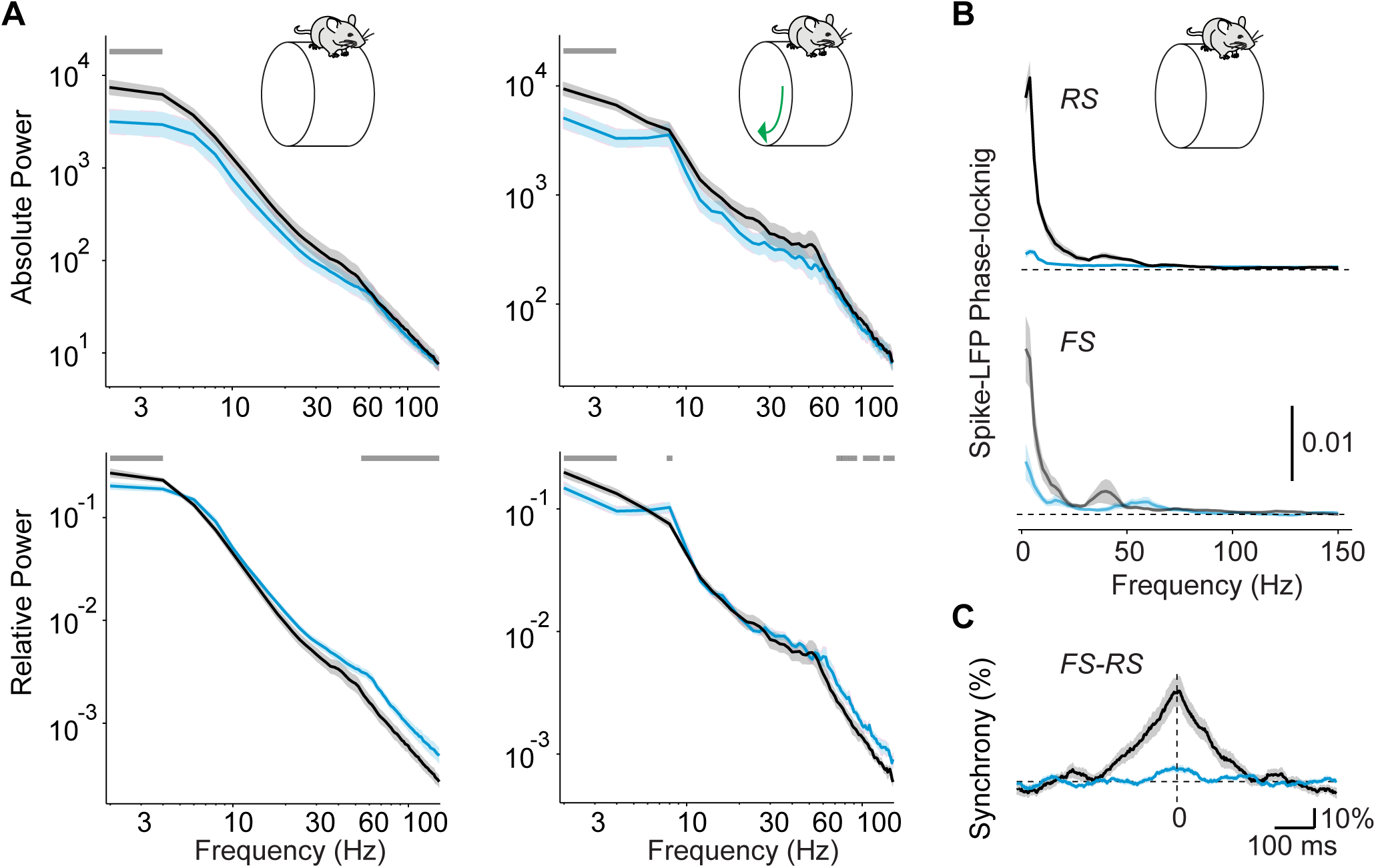
Power spectra from V1 recordings in ErbB4-VIP mutants and controls. (**A**) Upper, left: Absolute V1 LFP power for control (black) and mutant (cyan) populations during quiescence. Upper, right: Absolute LFP power spectra for control and mutant populations during locomotion, shown on a log-log scale. Lower, left: Relative LFP power spectra for control and mutant populations during quiescence. Relative power spectra were defined by dividing the power at each frequency by the summed power across frequencies. Lower, right: Relative LFP power spectra for control and mutant populations during locomotion. Controls: 21 sessions, 7 mice. Mutants: 29 sessions, 9 mice. The LFP is a complex signal that reflects thalamocortical and cortical inputs, as well as local RS and FS activity. The decrease in absolute power in low-frequency bands in mutants as compared to controls is consistent with a loss of synchrony (Figure 2), while the increase in relative high-frequency power is consistent with increased cortical firing rates in mutants (Einevoll et al 2013). Horizontal, gray bar at top indicates frequencies at which the difference between control and mutant mice was significant. (**B**) Average spike-LFP phase-locking (measured with pairwise phase consistency) during quiescence for RS (upper) and FS (lower) cells in controls (black) and mutants (cyan) (similar to Figure 2C). Both RS and FS cells in the ErbB4-VIP mutants showed decreased spike-LFP phase-locking at low- and gamma-range frequencies.. Controls: 55 RS, 23 FS cells, 7 mice. Mutants: 61 RS, 23 FS, 8 mice. (**C**) Average RS-FS cell synchrony during quiescence (similar to Figure 2E). Controls: n=15 pairs in 4 mice. RS-FS Mutants: n=95 pairs in 6 mice. Controls: n=85 pairs in 8 mice.

**Figure S6 (Related to Figure 3).**
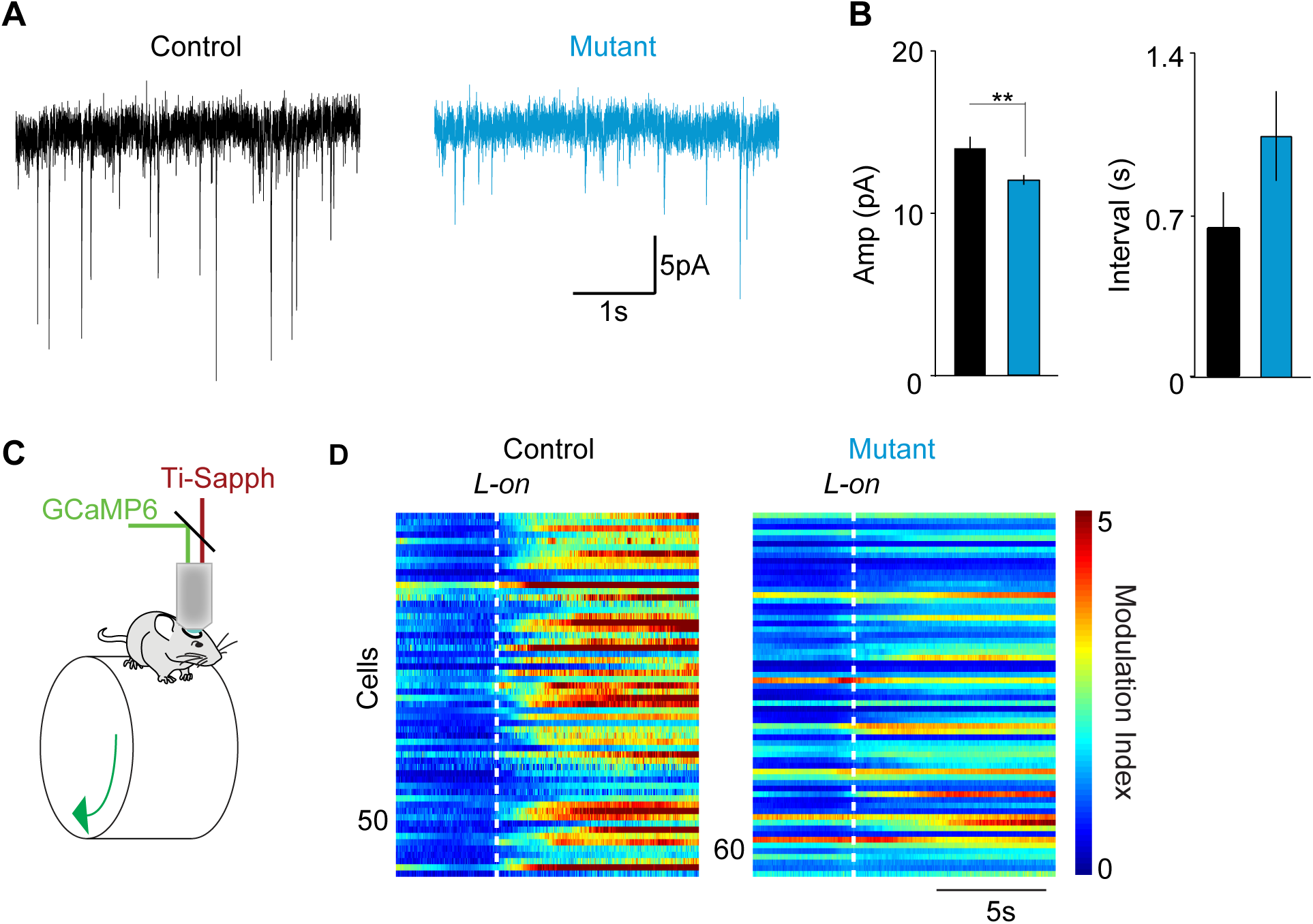
Deletion of ErbB4 reduces excitatory inputs to VIP interneurons and decreases their modulation by locomotion. (**A**) Example traces of miniature EPSCs (mEPSCs) recorded in VIP interneurons of control (left) and mutant mice. (**B**) Relative to controls (black), deletion of ErbB4 (cyan) results in a significant decrease in mEPSC amplitude (left) and a non-significant increase in inter-event interval (right). Controls: 6 cells, 4 mice. Mutants: 12 cells, 5 mice. (**C**) Schematic of the in vivo 2-photon imaging configuration. (**D**) Modulation of the activity of each VIP interneuron around locomotion onset in two example mice. **p<0.01

**Figure S7 (Related to Figure 5).**
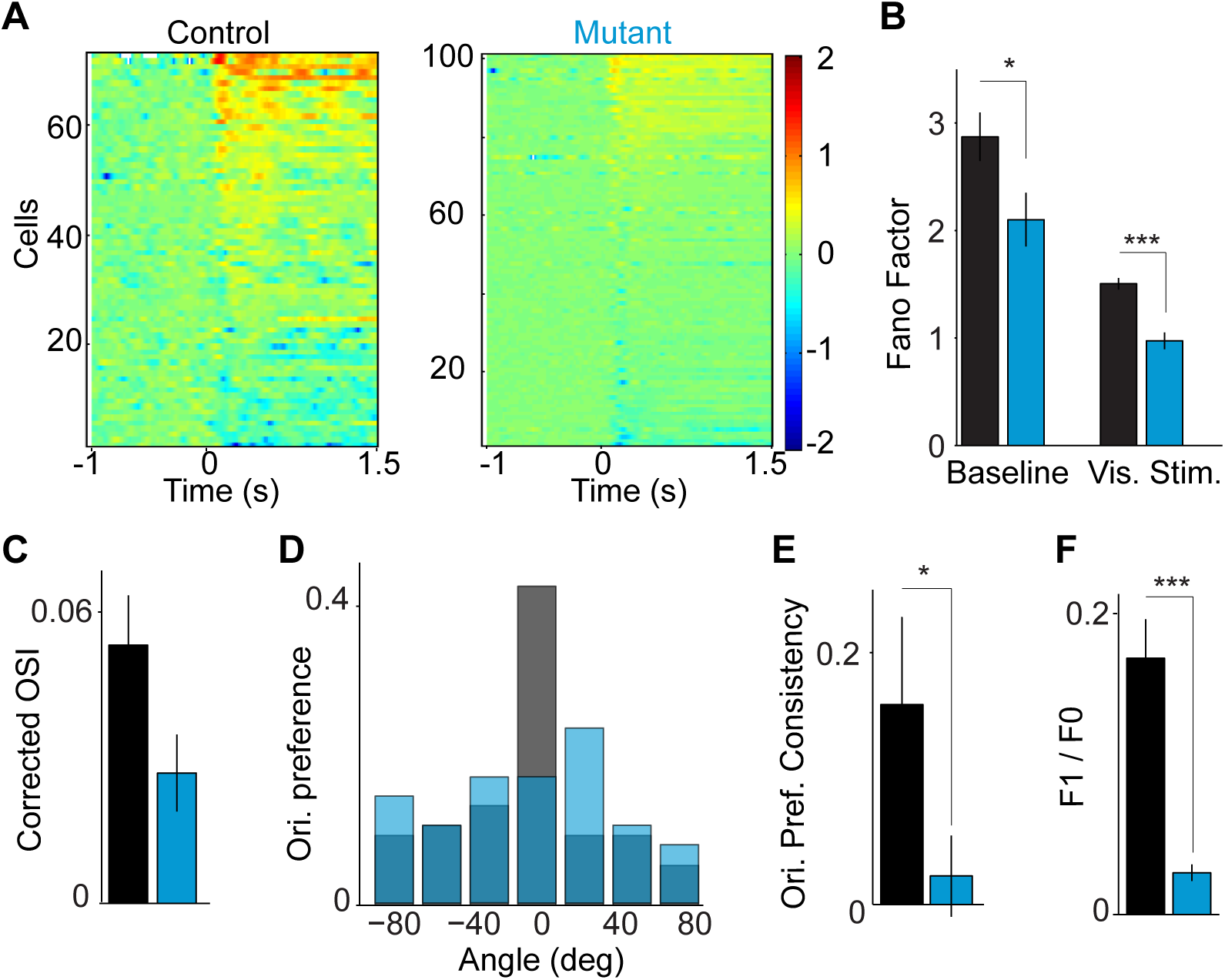
Additional quantification of visual response disruption in ErbB4-VIP mutants. (**A**) Shown for each individual RS cell the average rate modulation relative to the inter-trial interval baseline as a function of time around stimulus onset, i.e shown the rate modulation log10 FR(t) / FRbase. Left: control mice. Right: mutant mice. (**B**) Fano factor for the 30ms-500ms stimulus period and the inter-trial-interval (baseline) period. Fano factor was defined as the variance of the spike count divided by the mean of the spike count. (**C**) Average orientation selectivity index (OSI) with the random OSI that was obtained by shuffling trials subtracted. Corrected OSI values are higher than by chance for RS cells in both controls and mutants. (**D**) Histogram of preferred orientations across the population of RS cells for mutants and controls. (**E**) Population phase consistency of orientation preferences for controls and mutants. Controls exhibit significant clustering of preferred orientations, but mutants show no clustering. (**F**) Population average F1/F0 values for RS cells. Controls: 55 cells, 7 mice. Mutants: 61 cells, 8 mice. Error bars and shading show s.e.m.’s. p*<0.05, p**<0.01, p***<0.001.

**Figure S8 (Related to Figure 5).**
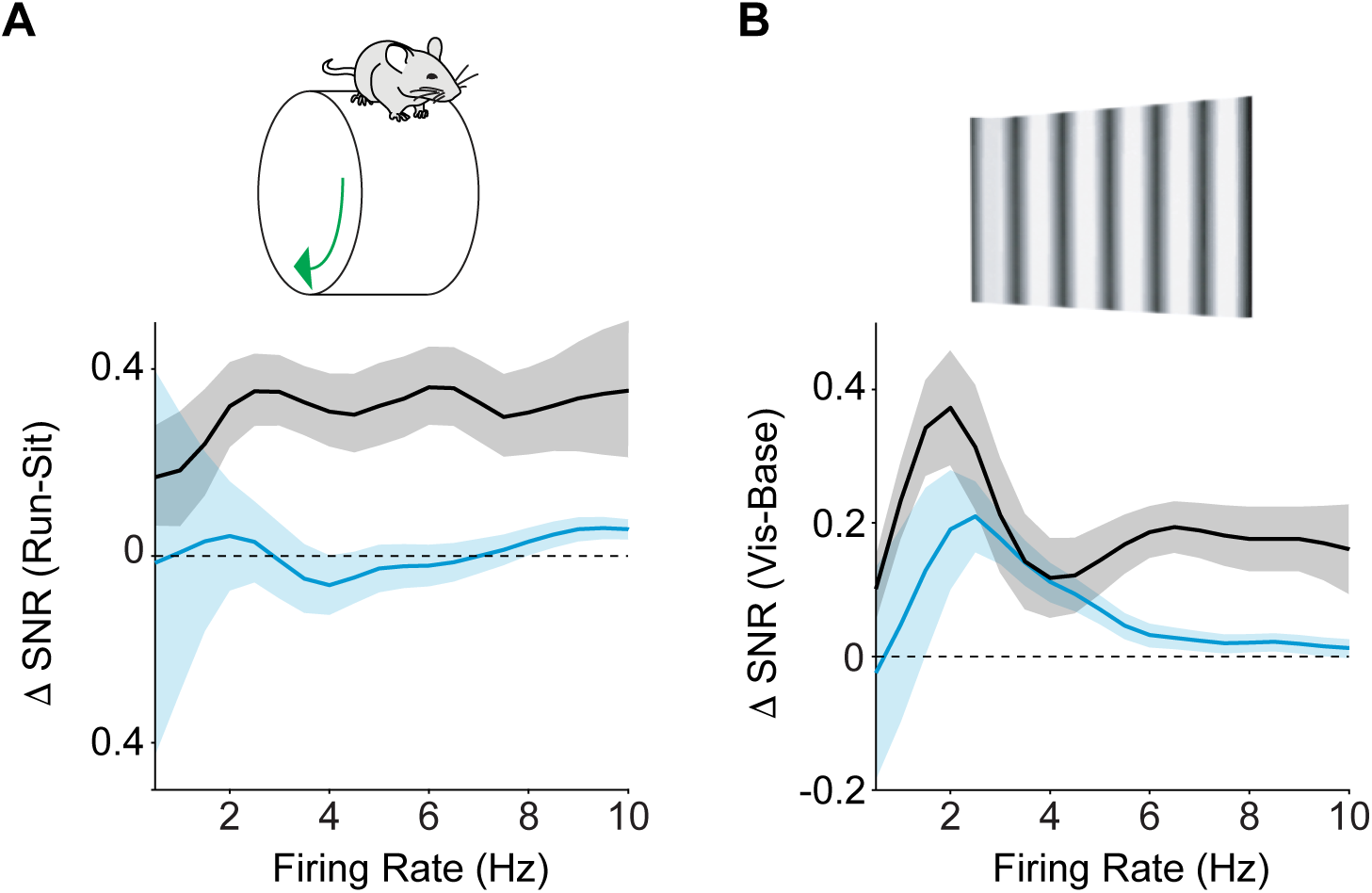
Effects of ErbB4-VIP deletion on state modulation and visual responsiveness are not due to a ceiling effect on firing rates. (**A**) Average rate modulation during early locomotion (-0.5 to 0.5s around L-on) period as compared to quiescence, as a function of firing rate for RS cells in controls and mutants.- Controls: 85 cells, 5 mice. Mutants: 72 cells, 8 mice. (**B**) Average rate modulation relative to inter-trial interval baseline as a function of time around stimulus onset. Controls: 106 cells, 8 mice. Mutants: 92 cells, 7 mice. If the differences in rate modulation between control and mutant mice were explained by a ceiling effect on RS firing rate, we would expect to find no differences in rate modulation for a given firing rate bin. However, RS cells in controls showed rate modulation even at higher firing rates. Shading shows s.e.m.’s.

**Figure S9 (Related to Figure 6).**
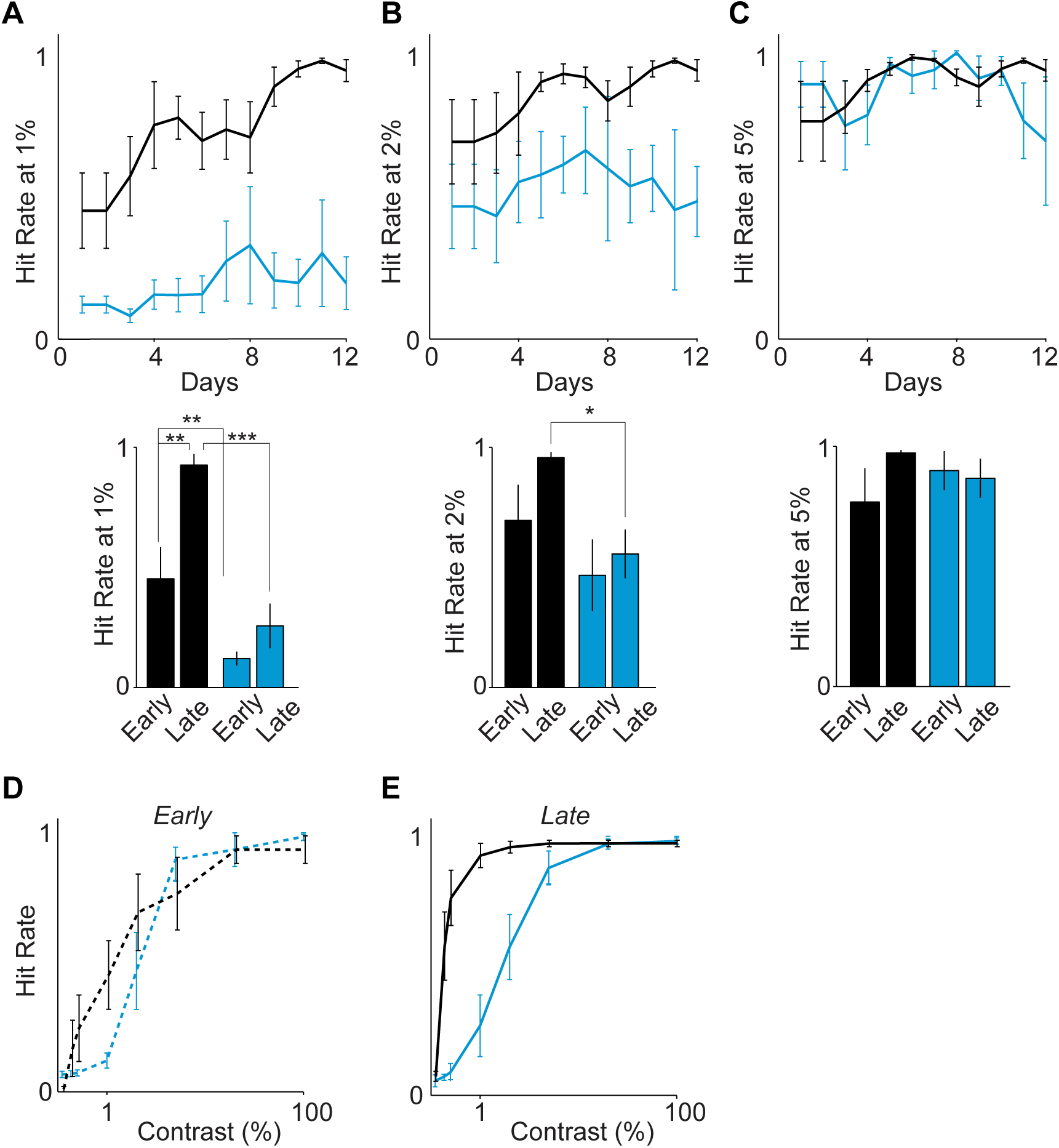
Visual task performance for control (black) and mutant (cyan) animals. (**A-C**) Top: Hit rates for 1% (**A**), 2% (**B**), and 5% (**C**) contrast stimuli over a period of two weeks. Bottom: Quantification of initial (Early) vs final (Late) performance of controls and mutants for each stimulus contrast level (similar to Figure 4K). (**D**) Psychophysical performance curves for both groups during the initial two training days. The initial psychophysical curves for mutants and controls were not significantly different except at the 1% level. (**E**) Performance curves for both groups during the final two training days. Whereas the controls showed learning, evident in the leftward shift of their performance curve between Early and Late training sessions, the mutant curve did not shift leftward over the two-week training period. Controls: 4 mice. Mutants: 4 mice. Error bars show s.e.m.’s. p*<0.05, p**<0.01, p***<0.001.

## SUPPLEMENTAL EXPERIMENTAL PROCEDURES

### Mouse Strains

The *Dlx6a*^*Cre*^ (Jackson, stock number 008199), *PV*^*Cre*^ (Jackson, stock number 008069), *VIP*^*Cre*^ (Jackson, stock number 010908), *SST*^*Cre*^ (Jackson, stock number 016868), *ErbB4*^*F/F*^ (Golub et al., 2004), and *Ai9* (Jackson, stock number 007905) reporter mouse lines were maintained on a C57/B16 background. Genotyping was performed as previously described. Crosses for histology included the *Ai9* reporter line. *Aci9* mice were crossed to *Dlx6a*^*Cre*^, *PV*^*Cre*^, or *SST*^*Cre*^, or *VIP*^*Cre*^ mice to generate *Dlx6a*^*Cre*^ Ai9, *PV*^*Cre*^ Ai9, or *SST*^*Cre*^ Ai9, or *VIP*^*Cre*^ Ai9 reporter animals.

### Data analysis

#### Quantification of firing rate

The firing rate was computed by dividing the total number of spikes a cell fired in a given period by the total duration of that period (Fig. 1E, F).

#### Quantification of inter-spike-interval maximum

For each cell, we computed the inter-spike-interval (ISI) histogram at 1ms resolution. We then determined the time at which this histogram had a peak. To obtain an average ISI histogram, we first normalized the ISI histogram of each cell individually by dividing by the maximum value in the ISI histogram (Fig. 1G).

#### Quantification of burstiness

We quantified the propensity to engage in burst firing using the coefficient of Local Variation (LV) (Shinomoto et al., 2009). The LV is a modification of the coefficient of variation (CV) measure and, like the CV, it quantifies spiking irregularity. A Poisson-process has an LV of 1. Rhythmic firing leads to LV values <1, whereas burst firing leads to LV values above 1. The LV quantifies irregularity only on the basis of pairs of subsequent ISIs, and, unlikely the CV, is therefore robust against non-stationarities in firing rate and the mean firing rate (Shinomoto et al., 2009). We also computed an alternative measure of burstiness, namely the log fraction of ISIs between 2-10 ms over the fraction of ISIs between 10 and 100 ms, log(ISI_short_ / ISI_long_). We have previously shown that there is a strong correlation between the latter measure and the LV (Vinck et al., 2015).

#### Computation of wheel position and change points

Wheel position was extracted from the output of the angle sensor. Since wheel position is a circular variable, we first transformed the sensor data to the [-π,π] interval. Because the position data would make sudden jumps around values of π and –π, we further performed circular unwrapping of the position phases to create a linear variable (Fig. 2A).

We then used a change-point detection algorithm that detected statistical differences in the distribution of locomotion velocities across time. The motivation of this method relative to the standard method of using an arbitrary threshold (e.g., 1 cm/s) (Niell and Stryker, 2010) is that our technique allowed for small perturbations in locomotion speed to be identified that might otherwise fail to reach the locomotion threshold. Further, it ensured that the onset of locomotion could be detected before the speed reached 1 cm/s. If the distributions of data points 100 ms before and 100 ms after a certain time point *t* were significantly different from each other, using a standard T-test at p<0.05 and sampling at 2 kHz, then the data point was deemed a candidate change point. A point *t* was considered a candidate locomotion onset point if the speed 100 ms after *t* was significantly higher than 100 ms before *t*. A point was considered a candidate locomotion offset point if the speed 100 ms after *t* was significantly lower than 100 ms before *t*. A point was accepted as a locomotion onset point if the previous transition point was a locomotion offset point. A point was considered to be a locomotion offset if it was preceded by a locomotion onset point, and if the speed 100 ms after *t* did not significantly differ from zero. This prevented a decrease in speed to be identified as a locomotion offset point. We further required that a locomotion offset point not be followed by a locomotion onset point for at least 2 s, because mice sometimes showed brief interruptions between bouts of running.

We selected locomotion trials for which the average speed until the next locomotion offset point exceeded 1 cm/s and which lasted longer than 2 s. Quiescence trials were selected that lasted longer than 5 s, had an average speed <1 cm/s, and for which the maximum range of movement was <3 cm across the complete quiescence trial.

#### LFP power

To compute LFP power spectra, we divided the data in 500 ms periods and multiplied each data segment by a Hann taper. We then computed the average LFP power spectrum by computing the FFT per segment and averaging over the segment’s power spectra.

#### Spike-field locking analysis

Spike-field locking was computed using the Pairwise Phase Consistency (Vinck et al., 2012), a measure of phase consistency that is not biased by the firing rate or the number of spikes. The PPC is computed as follows:

1. For each spike, a spike-LFP phase is computed at each frequency (see below).
2. For each pair of spikes (fired by one cell) that fell in a different trial, we then compute the inner product of the two spike-LFP phases (the inner product being a measure of their similarity). Spike-LFP phases were computed for each spike and frequency separately by using Discrete Fourier Transform with Hanning taper of an LFP segment of length 9/*f*, where *f* is the frequency of interest. For a given time period (quiescence or locomotion), we only selected cells that fired at least 50 spikes in that period.
3. The PPC then equals the average of the inner products across all pairs of spike-LFP phases that fell in different trials. Note that exclusively taking pairs of spike-LFP phases from different trials ensures that history effects like bursting do not artificially inflate the measure of phase locking. The expected value of the PPC ranges between 0 and 1, although estimates lower than zero can occur.

For each cell, we computed the preferred phase of firing in the 40-60 Hz range by first computing the circular mean across all spike-LFP phases, and then computing the circular mean across frequencies (Fig. 2D, left).

We then computed the phase consistency (Fig. 2D, right) of preferred spike-LFP phases across units by computing the PPC over the preferred spike-LFP phases. We then computed an estimate of the standard error of the mean using the jack-knife (Fig. 2D, right).

#### Computation of STAs

To compute the average STA in the delta/theta [1,6] Hz and the gamma [40-60] Hz band, we first filtered the LFP data of 2s traces around each spike. We then normalized the energy of the LFP trace by either dividing by the mean absolute value of the LFP signal. We then averaged these traces across spikes.

#### Computation of pair correlations

Unit-unit correlations (Fig. 2E) were computed using the cross-correlogram at 1ms resolution. The cross-correlogram contains, for each cell pair, the number of spike coincidences at a certain delay (e.g. the number of times one neuron fired a spike 5-6 ms after another neuron). We normalized the cross-correlogram by computing the percent-wise increase compared to the expected fraction of coincidences given the firing rates of the two cells.

#### Computation of modulation by state

To examine whether RS and FS cell firing rates were significantly changed around locomotion onset, we computed the firing rate in the [-0.5, 0.5] s window around locomotion onset (L-on; as in (Vinck et al., 2015) and compared this to the firing rate in the [-5,-2] s quiescence period before locomotion onset by computing log(FR_L-on_ / FR_Q_) (Fig. 3C, Fig. 4C).

Using the same modulation index, we also compared the firing rate in the early [2,5] s quiescence period after locomotion offset – when the animal was still aroused but not yet moving (Vinck et al., 2015) – to the late [>40] s quiescence period after locomotion onset, when the animal had low arousal levels and is not moving (Vinck et al., 2015) (Fig. 3D, Fig. 4D).

#### Computation calcium signals

Analysis of imaging data was performed using ImageJ and custom routines in MATLAB (The Mathworks). Motion artifacts and drifts in the Ca^2+^ signal were corrected with the moco plug-in in ImageJ (Dubbs et al., 2016), and regions of interest (ROIs) were selected as previously described (Chen et al., 2013). All pixels in a given ROI were averaged as a measure of fluorescence, and the neuropil signal was subtracted.

#### Quantification calcium signals

Frame times from the resonant scanner were used to align the Ca^2+^ signals with the wheel traces. Ca^2+^ signals were expressed as ΔF/F(t). Briefly, a time-dependent baseline, F0(t), was taken as the average of the minimum 10% of overall fluorescence, F(t), during the recording period. The relative change in fluorescence was calculated as ΔF/F(t)=(F(t)- F0(t))/F0(t). For this analysis, we selected locomotion trials which lasted 5s or longer, and quiescent trials which lasted 15s or longer. To determine whether Ca^2+^ activity was altered during behavioral state transitions, ΔF/F(t) from [0,4]s after locomotion onset (Ca_L-on_) was compared with ΔF/F(t) from [10,15]s after locomotion offset (Ca_Q_) by computing (Ca_L-on_ - Ca_Q_)/(Ca_L-on_ +Ca_Q_).

#### In vitro electrophysiology analysis

Custom-written algorithms in Igor Pro (M.J. Higley; Wavemetrics) were used to detect and measure miniature events based on a template-matching method (Clements and Bekkers, 1997). Data were compared using Student’s t-tests.

#### Quantification rate modulation to visual stimulus

For this analysis we computed the firing rate in the 30-500 ms period after stimulus onset, and the firing rate in the 1000 ms before stimulus onset. We then computed the firing rate modulation of stimulus-driven versus baseline rate as log(FR_stim_ / FR_base_) (Fig. 5D). We call this modulation the SNR. We also computed the SNR at each time point (Fig. 5B, C) by computing log(FR_stim_(t) / FR_base_) where FR_stim_(t) is the estimated firing rate at time *t*, which was estimated by convolving the spike trains with Gaussian smoothing windows (50ms, sigma=12.5ms), We further compared this modulation between the entire locomotion period and the entire quiescence period, computing the ΔSNR as log(FR_stim, L_ / FR_base, L_) - log(FR_stim, Q_ / FR_base, Q_) (Fig. 5G).

#### Computation of modulation (F1/F0)

To determine the extent to which cells showed linear responses to a drifting grating, we performed the following analysis. We first computed the average spike density of the firing by convolving the spike trains with Gaussian smoothing windows (50ms, sigma=12.5ms), and averaging these over trials. We then computed a Fourier Transform of the average spike density in the 0.25-1.25 s window, thereby excluding the initial firing transient. (Note that including this transient would likely lead to artificially increased F1/F0 values). The F1 component was extracted as 2 times the amplitude of the Fourier component at 2 Hz (the temporal frequency of the drifting grating), and the F0 component was taken as the 0 Hz, DC component of the FFT. We then computed an index of linearity as L=F1/F0.

#### Quantification of orientation selectivity

The orientation selectivity index is defined as R = [1-Circular Variance]. This is derived by letting each orientation (measured in radians, with 0 and 90 degree orientations corresponding to 0 and PI radians) be a vector on the circle with weight 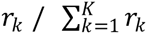. Note that we performed this procedure for the directions lying between 0 and 180 degrees and 180 and 360 degrees separately, and averaged all derived measures over the two set of directions. We then computed the resultant vector by summing the sine and cosine components.

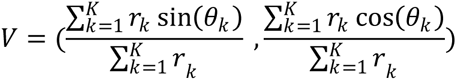

We next computed the resultant vector length (OSI) as R = |V|. Note that 0<=R<=1, and that R=1 indicates that a cell only has a non-zero firing rate for one orientation, whereas R=0 indicates that the cell has the same firing rates for all orientations. These vectors can also be normalized to unity length by V/|V|. These vectors are shown on the unit circle in Fig. 5F, right, and shown in histogram form in Fig. S7E. We also computed the phase consistency over the preferred orientations similar to Fig. S7D.

The OSI is a biased quantity in that it tends to be over-estimated for a finite number of trials (Womelsdorf et al., 2012). This can be intuitively seen: Suppose that R=0, meaning that the true firing rate for each orientation is identical. With a small number of trials, the firing rate estimates will deviate from those true estimates, which causes R>0. This bias is typically also stronger if the firing rate is lower (Womelsdorf et al., 2012). The reason for that is that the ratio mean / s.e.m. (where s.e.m. is the standard error of the mean) tends to be higher when the firing rate is high, assuming a Poisson process. That is, proportionally speaking, the deviation in the estimated firing rate from the true firing rate tends to be larger when the true firing rate is low.

To correct for these intrinsic biases, we performed the following estimation procedures. 1) For each cell, we randomly selected 50 spikes across all spikes and then compute the OSI. We repeated this procedure by bootstrapping a random sample of 50 spikes (without replacement) for 10000 times and computing the average OSI across these bootstraps. These estimates are shown in Fig. 5F. 2) For each cell, we randomly shuffled the trials across orientations. This was done such that if the first orientation originally contained 10 trials, the surrogate/random set of trials for the first orientation would also contain 10 trials. We then computed the OSI for the shuffled condition, and repeated this procedure 5000 times. In order to correct for the bias, we then subtracted the average shuffled OSI from the true OSI. These values are shown in Fig. S7C. Values greater than zero indicate that there was more orientation than by chance.

#### Computation of correlations between FR and SNR

To compute the correlation between FR and the rate modulation by visual stimulus, we computed the average firing rate over baseline and stimulus periods as FR_avg_ = (FR_stim_ + FR_base_)/2. We then binned FR_avg_ by constructing bins that were B = [bincenter-0.2*bincenter, bincenter+0.2*bincenter] wide. For each bin, we then computed the average rate modulation by the visual stimulus.

We performed the same procedure for the modulation by state. In this case FR_avg_ was computed as FR_avg_ = (FR_Q_ + FR_L-ON_)/2.

#### Computation of visual performance false alarm rate

The False Alarm Rate was defined as the average number of incorrect hits, i.e. licking responses when the GO stimulus was not displayed (i.e. the NOGO condition).

#### Computation of visual performance curves

For each session, we corrected the total number of true hits by the false alarm rate as follows. Assuming that each time when the mouse sees the stimulus, the mouse responds to the stimulus, observed HITRATE equals HITRATE_observed_ = HITRATE_true_ + (1- HITRATE_true_)*FAR.

Thus, it follows that

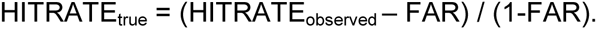

If HITRATE_observed_ < FAR, then we let HITRATE_true_ = 0. If the observed hit rate is 100%, then the true hit rate is also 100%. If the observed hit rate equals the FAR or is smaller than the FAR, then the estimated true hitrate equals zero.

We cleaned up the data per session automatically as follows. First, we ensured that when the mice stopped performing at the end of a session, this data was not incorporated into the average. This was done by computing a 10-point running moving average of the data. For the *k*-th trial, we then computed the average performance of the mouse (as HITRATE_observed_) until the (k-1)-th trial. This average performance was computed starting from the trial where the mouse had obtained at least 10 rewards, to prevent poor performance at the start from influencing the average. (Note that the first ten trials in a given session were always 100% contrast trials). The same procedure was performed on the FAR. The last trial was defined as the trial at which the 10-point moving average of the hit rate or the false alarm rate fell below 75% of the mean performance up to that point and did not recover above this level anymore.

We then computed, for each contrast, the average hit rate for each contrast and the FAR, and estimated the true hit rate as detailed above. We fit sigmoid curves to the contrast vs. hit-rate data. We constructed learning curves by computing the 2-day moving averages of the psychophysics curve for each mouse. We then took the value of the 2-day moving average at a given contrast and computed the mean and s.e.m. across mice. These values are shown in Fig. 6D. We also computed the average psychophysics curve for the first two sessions, and for all the sessions starting from day 12, when the performance of the controls reached a saturation point. These curves are shown, at the measured contrasts, in Fig. 6B. We performed a standard T-test to test for differences between animals, and to test for differences between early and late trials.

#### Statistical testing

A common problem in many experimental studies is the use of nested design, where multiple cells are measured for each animal and cannot be taken as independent measurements (Galbraith et al., 2010). Thus, directly performing statistical comparisons between a sample of control and mutant cells, as is typical in the neurosciences, leads to a highly increased false alarm rate and suboptimal estimates of the mean (Aarts et al., 2014).

To avoid the increased false positive rate inherent in nested designs, we used semi-weighted error estimators, commonly used in random-effects meta-analysis (Chung et al., 2013; DerSimonian and Laird, 1986). Let y_i_ be the mean for one parameter (e.g., OSI) for the *i-*th animal, where x_j_ is the parameter value for the j-th cell and there are M cells per animal, defined as:

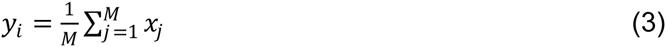

The unweighted estimator of the mean is defined as the mean over y_i_, using the number of animals per condition as degrees of freedom. This analysis is suboptimal, as some animals have many cells and yield more reliable estimates, whereas other animals have few cells and yield more unreliable estimates. Alternatively, a weighted estimator can be defined by pooling observations across N animals and using the number of cells per condition as degrees of freedom. Although this estimator is commonly used in the neurosciences, statistical inference based on pooling cells together does not properly control the false alarm rate (Aarts et al., 2014)

The semi-weighted estimator, for a given experimental condition (e.g., CT cells) is defined as:

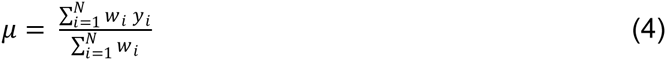

where weight for the i-th animal is defined as:

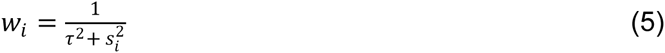

Here, 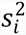 is the within-animal variance across cells and τ^2^ is the estimated variance across all animals of a given condition using the maximum likelihood method (Chung et al., 2013):

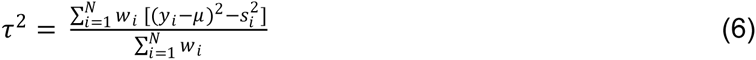

We solved equations 5 and 6 iteratively and checked for convergence of τ^2^ to <0.001. Intuitively, if there is no across-animal variance, then the contribution of each cell is weighted by the number of animals (the weighted estimator). If the across animal variance is very large (i.e., observations within an animal are maximally dependent), then each animal is given the same weight. Previously, we used this semi-weighted estimator to calculate the statistical significance of the difference between cell populations using a standard Student’s t-test (Lur et al., 2016).

For this paper, we improved on this previous statistical procedure by developing a nonparametric statistical inference procedure to test for statistical differences between animals. Using a nonparametric statistical permutation test avoids any assumptions inherent to parametric statistics. However, directly permuting animals between controls and knockouts would lead to a test with low statistical sensitivity because it can yield permutations in which one group could contain many animals with a low number of cells (which would yield a large variance of the permutation distribution). We therefore used a stratified permutation test that circumvents this problem and that was constructed as follows:

We ranked the animals according to the number of cells recorded (which is roughly inversely proportional to the variance). We then created ‘strata’ containing, for each condition, a subset of animals that had similar ranks. This was done as follows. For the condition with the fewest number of animals, we ranked the animals according to the number of cells they had and ranked them into non-overlapping strata of 2. If the number of animals was uneven then the last strata would contain 3 animals. We then created the same number of strata for the other condition and placed them in these strata according to their ranks.

For example, the division of strata for the following distributions of animal and cell numbers would be as follows

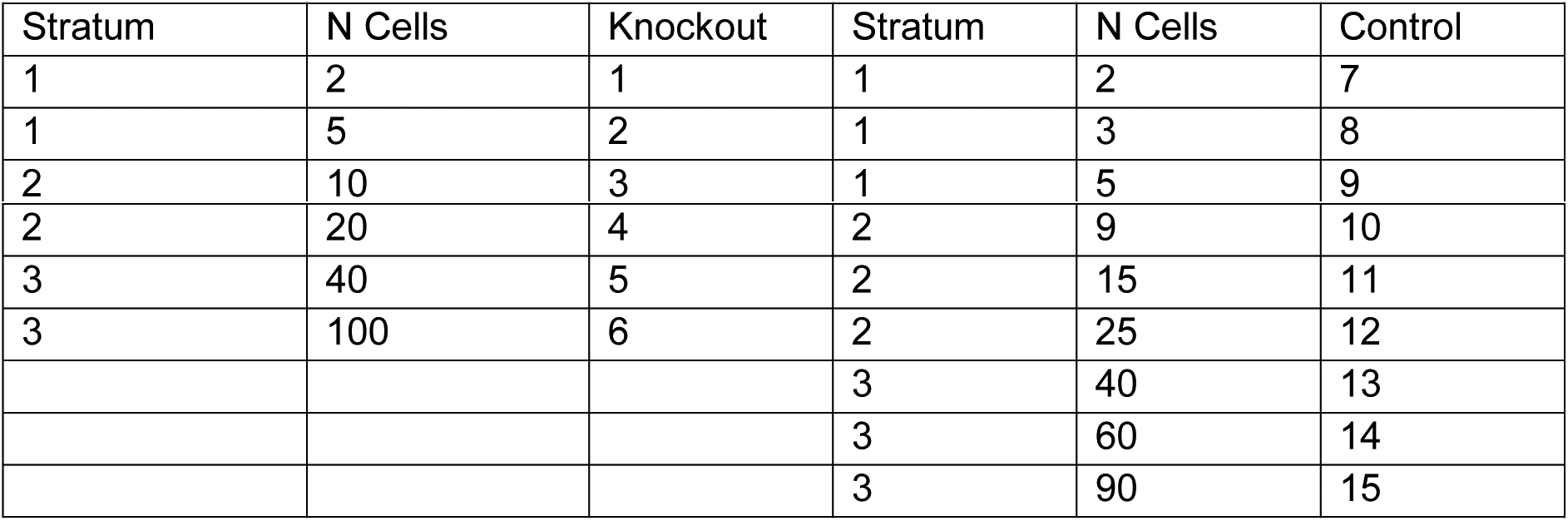

| Stratum | N Cells | Knockout | Stratum | N Cells | Control |
| --- | --- | --- | --- | --- | --- |
| 1 | 2 | 1 | 1 | 2 | 7 |
| 1 | 5 | 2 | 1 | 3 | 8 |
| 2 | 10 | 3 | 1 | 5 | 9 |
| 2 | 20 | 4 | 2 | 9 | 10 |
| 3 | 40 | 5 | 2 | 15 | 11 |
| 3 | 100 | 6 | 2 | 25 | 12 |
|  |  |  | 3 | 40 | 13 |
|  |  |  | 3 | 60 | 14 |
|  |  |  | 3 | 90 | 15 |

Then, one permutation would look as follows

| Stratum | N Cells | Random 1 | Stratum | N Cells | Random 2 |
| --- | --- | --- | --- | --- | --- |
| 1 | 2 | 1 | 1 | 5 | 2 |
| 1 | 2 | 7 | 1 | 3 | 8 |
| 2 | 10 | 3 | 1 | 5 | 9 |
| 2 | 9 | 10 | 2 | 20 | 4 |
| 3 | 40 | 13 | 2 | 15 | 11 |
| 3 | 60 | 14 | 2 | 25 | 12 |
|  |  |  | 3 | 40 | 5 |
|  |  |  | 3 | 100 | 6 |
|  |  |  | 3 | 90 | 15 |

We then computed, for each condition, the semiweighted estimator, which yields a difference between the semiweighted estimator in the control and the mutant condition. For each random permutation, we also computed the difference between the semiweighted estimator for the random group 1 and the random group 2. This yielded a randomization distribution of differences between the conditions. Under the null hypothesis, the data are exchangeable between the knockout and the control conditions (i.e. their statistical distributions are identical). We tested against this null hypothesis by comparing the observed difference in semiweighted estimators with the 95% percentile of the randomization distribution.

We compared the resulting p-values with comparing the mean and sem values of the semi-weighted estimators according to the T-distribution (as in (Lur et al., 2016)), which generally agreed very well. Compared to directly testing for differences between cells as is commonly done in neuroscience, the p-values we observed were often a few orders of magnitude larger, while our procedure was generally more sensitive than computing the mean per animal first and performing statistics over those per-animal means.

At all places in the manuscript, we used this stratified permutation test over the semi-weighted estimators to test for statistical differences. In the figures where we show pairwise differences between conditions, for example locomotion vs. quiescence and stimulus vs. baseline, we show weighted means (Fig. 3C, Fig. 5D, G). In those figures where we report spike-spike and spike-field correlations, we also show weighted means. This allows one to judge whether these differences and correlations are significantly different from zero. In these cases, we always performed statistical testing over the semi-weighted means of the pairwise differences. For other figures where this is not relevant, for example absolute firing rates or LV values, we show the semi-weighted means.

